# Thalamus drives active dendritic computations in cortex

**DOI:** 10.1101/2021.10.21.465325

**Authors:** Arco Bast, Jason M. Guest, Rieke Fruengel, Rajeevan T. Narayanan, Christiaan P.J. de Kock, Marcel Oberlaender

## Abstract

Perception is linked to a calcium-dependent dendritic spiking mechanism that enables the major output cells of the cerebral cortex – layer 5 pyramidal tract neurons – to combine inputs from different information streams. Which circuits activate this mechanism upon sensory input is unclear. Here we found that thalamocortical axons, which provide sensory input to cortex, target specifically the dendritic domains in pyramidal tract neurons that initiate calcium spikes. Sensory input thereby enables distal dendritic inputs preceding the stimulus to transform the first responses that leave cortex into bursts of action potentials. Thus, thalamus can drive active dendritic coupling of sensory with prestimulus information streams to modulate cortical output. Our findings indicate that thalamocortical coupling is first in a cascade of mechanisms that transform sensory input into perception.

## Introduction

The ability to combine sensory signals from the environment with internally generated information streams is a hallmark feature of the cerebral cortex, forming the basis for perception and cognition, and the resulting behavior. It is thought that the pyramidal tract neurons in cortical layer 5 (L5PTs) are key for such combination processes (*1, 2*). Along extensive dendrites, these major cortical output cells combine information streams arriving at all layers, and then broadcast the results of this combination to a variety of subcortical brain regions (*3*). Distal dendritic inputs, which typically appear attenuated at the soma, must cross a high threshold to evoke dendritic calcium action potentials (APs) (*4*), but can then modulate cortical output with bursts of APs (*5*). Evidence that this *in vitro* observed calcium AP-dependent burst firing mechanism is utilized to implement higher brain functions was recently reported for L5PTs in the vibrissa-related part of mouse primary somatosensory cortex (vS1) – the barrel cortex (*6, 7*). Elevated calcium activity in distal dendrites in conjunction with sensory-evoked burst firing correlated with the threshold for perceiving whisker stimuli. Manipulating the dendritic calcium domain even allowed to lower or raise the perceptual threshold. The circuits underlying these remarkable observations have yet to be revealed. This is challenging, as it requires dissecting of how synaptic inputs from different thalamocortical (TC), intracortical (IC) and top-down corticocortical (CC) pathways contribute to sensory responses in neurons with complex dendritic physiology.

Thalamus is the main route by which sensory signals reach cortex. Axons from primary thalamic nuclei terminate most densely in layer 4, from where sensory signals then propagate along the canonical L4→L2/3→L5/6 pathway (*8-10*). However, it has become increasingly clear that L5PT responses do not rely on serial signal flow through the canonical pathway. The onsets of L5PT responses rival or even precede those in layer 4 (*11-15*), and inactivation of layer 4 does not abolish L5PT responses (*12*). It was hence suggested that L5PTs are driven directly by thalamus (*12*). However, we recently found that this is not the case. Instead, we showed that L5PTs are indirectly driven by thalamus via corticocortical neurons (L6CCs) that cluster around the second innervation peak of primary TC axons at the layer 5 to 6 border (*14*). These L6CCs show the fastest and most reliable sensory responses in cortex, have elaborate axons within layer 5, and their inactivation abolishes L5PT responses (*14*). The TC axons that drive responses of L6CCs and in layer 4 (*16*), also converge strongly onto L5PTs (*12*). This direct input from thalamus extends across both the basal and apical dendrites of L5PTs (*17, 18*), including the dendritic domains that initiate calcium APs. Therefore, here we asked: *What is the function of the TC→L5PT pathway, and could this direct sensory input to the major cortical output cells contribute to their role during perception?*

To address these questions quantitatively, we combined electrophysiological measurements and neuroanatomical reconstructions in the thalamocortical whisker system of the rat with multi-scale modelling and optogenetic manipulations. Our multidisciplinary approach provided several lines of evidence that reveal how direct sensory input from thalamus contributes to responses in L5PTs – and in particular to burst firing. We found that TC synapses accumulate most densely around the dendritic domains that initiate calcium APs in L5PTs. We show that L5PTs elicit bursts of APs that precede responses in layer 4 upon both passive stimulation in anesthetized animals and during active sensing in awake animals. We demonstrate across a wide range of model configurations that L5PTs require direct TC input to generate such fast burst responses, that fast burst responses with 3 APs provide a readout for TC-driven calcium APs, and that prestimulus depolarization by distal inputs facilitates the occurrence of TC-driven calcium APs. Finally, optogenetic manipulations of *posthoc* reconstructed and simulated L5PTs provide *in vivo* evidence in support of the *in silico* predicted function of the TC→L5PT pathway – demonstrating that cortical output transitions from single APs to bursts with 3 APs depending on the amount of prestimulus inputs by IC neurons.

## Results

We quantified the distribution of TC synapses along the dendrites of L5PTs **(Fig. 1a-c)**. For this purpose, we injected an adeno-associated virus (AAV) into the primary thalamus of the whisker system **(Fig. S1a-b)** – the ventral posterior medial nucleus (VPM). In virus injected rats, we used cell-attached recording pipettes to *in vivo* label individual neurons in barrel cortex for *posthoc* reconstruction and cell type classification **(Fig. S1c)**. We identified TC synapses based on super-resolution light microscopy, if the vesicular glutamate transporter 2 (VGlut2) – a marker for TC synapses **(Fig. 1a-b)** – was expressed at contact sites between dendritic spines and virus-infected axonal boutons (13,296 contacts, 10 L5PTs, median VGlut2-positive contacts: 86%). The majority of TC synapses was distributed along the proximal (i.e., basal and apical oblique) dendrites **(Fig. 1d)**. However, synapse densities were highest along the apical trunk **(Fig. 1d-e)**. Surprisingly, for each L5PT, we observed a density peak close to the primary branch point (BP) of the apical trunk, irrespective of its distance to the soma **(Fig. 1e-f)**. Thus, TC synapse densities gradually increase with soma distance along the apical trunk, reach a peak ∼100 μm below the primary branch point, and then decay exponentially towards the apical tufts **(Fig. 1g)**. This highly specific innervation of the apical dendrites by primary thalamus cannot be explained by axo-dendritic overlap **(Fig. S1d)**, or by higher spine densities **(Fig. S1e)**, and it was not apparent in other cell types **(Fig. S1f)**.

**Fig. 1:**
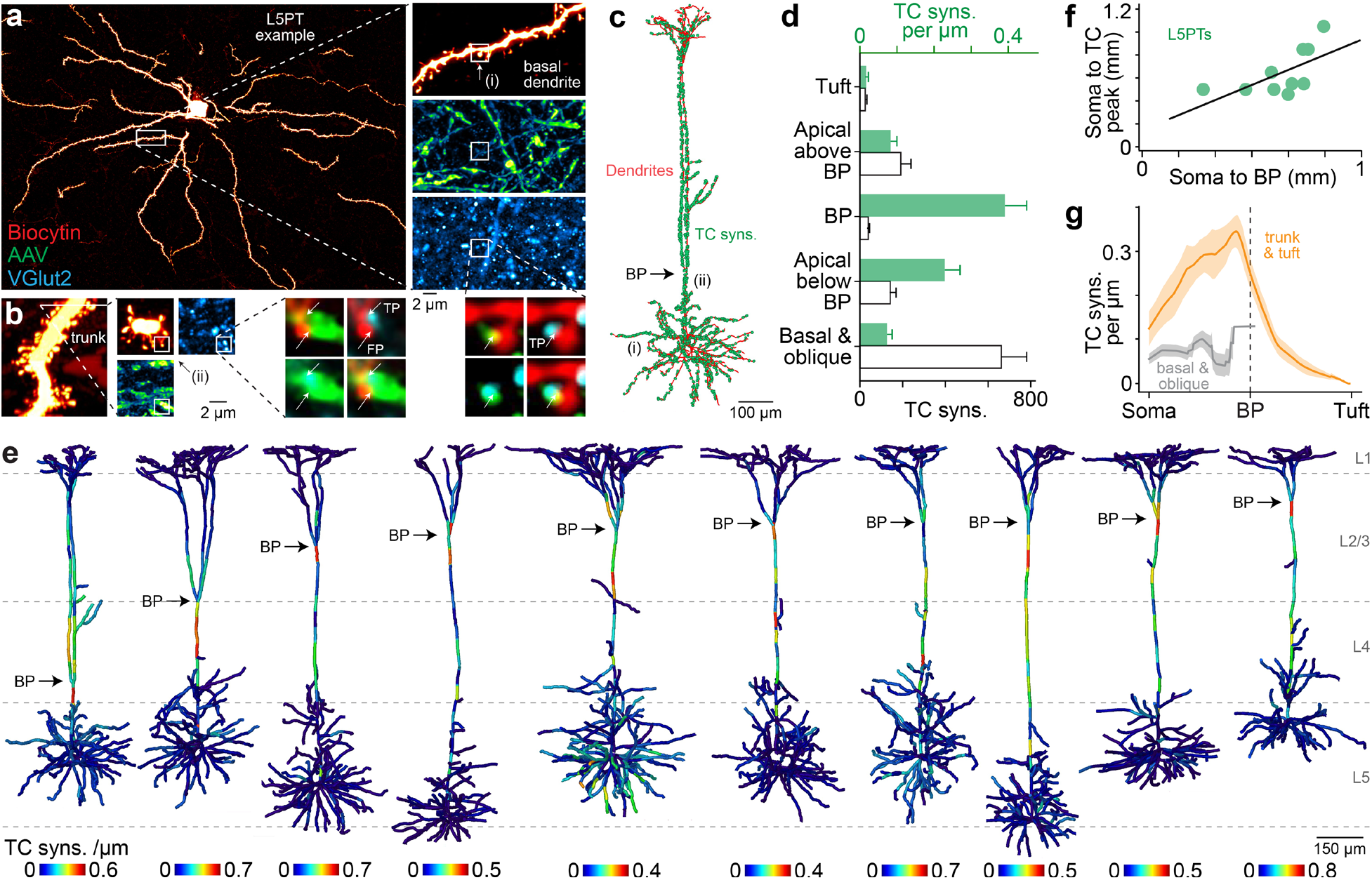
Anatomy of TC→L5PT pathway indicates contribution to sensory-evoked bursts. **a**. Left: example of biocytin labeled L5PT in vS1. Right: super-resolution images of spines along basal dendrite (top), boutons along TC axons (center) infected by AVV injections **(Fig. S1a**), and VGlut2 puncta (bottom). Zoom-ins: we identify ‘true positive’ (TP) synapses (arrow) if VGlut2 is expressed (cyan) at contact sites between spine heads (red) and TC boutons (green). **b**. Left: apical trunk of the L5PT from panel a. We show a side view that we generated by digitally resectioning the super-resolution image stack. The white line denotes the optical section for the zoom-ins (center) in which we show biocytin, AAV and VGlut2 labeling as top views. Right: zoom-ins show example TC bouton that contacts two spine heads (arrows). Only one of the contact sites expresses VGlut2, and we hence identify them as TP and ‘false positive’ (FP) TC synapses, respectively. **c**. 3D reconstruction of the dendrites (red) and TC synapses (green) of the L5PT shown in panel a. (i) and (ii) denote the respective depths of the TC synapses from panel a. **d**. Number (black) and density (green) of TC synapses along different dendritic compartments (mean ± STD). **e**. Density of TC synapses along L5PT dendrites. **f**. The BP location correlates significantly with the location of highest TC synapse density (Pearson R=0.8, p<0.01). **g**. We aligned L5PTs by their BP and plotted TC synapse densities (mean ± SEM) along basal and apical oblique dendrites (grey), and apical trunk and tuft dendrites (orange).

The wiring specificity in the TC→L5PT pathway resembles calcium (Ca^2+^) influx into the apical dendrites, which also increases towards a peak near the primary branch point (*19*). It is therefore likely that TC input facilitates the initiation of calcium APs (*4, 19*), and hence directly contributes to sensory-evoked bursts in L5PTs (*5, 20, 21*). To test this possibility, we recorded the activity of L5PTs (n=25 including those with reconstructed TC synapses) in the barrel cortex of anesthetized rats. L5PTs responded reliably to the onsets of passive whisker deflections **(Fig. 2a)**. Within the same L5PTs, responses varied across trials between three types: single APs, bursts with 2 or 3 APs **(Fig. 2b)**. Bursts increased significantly in L5PTs (Wilcoxon signed-rank: p<0.02) upon stimulus onset **(Fig. 2c)**, preceded responses of layer 4 spiny neurons (L4SP), but succeeded those of L6CCs (ANOVA with multiple comparison: p<0.05 for L4SP, p<0.01 for L6CC). In line with these cell type-specific response latencies, local field potential (LFP) recordings demonstrated that TC input to layers 5/6 and 3/4 precede bursts in L5PTs (p<0.01), on average by 3.4 and 1.6 ms, respectively **(Fig. 2d)**. Thus, our anatomy and physiology data indicate that thalamus could contribute to burst responses in L5PTs in three ways: indirectly via L6CCs (*14*), directly via the low density but high number of TC synapses along proximal dendrites, and directly via the high density of TC synapses along the apical trunk – particularly around the calcium domain near the primary branch point.

**Fig. 2:**
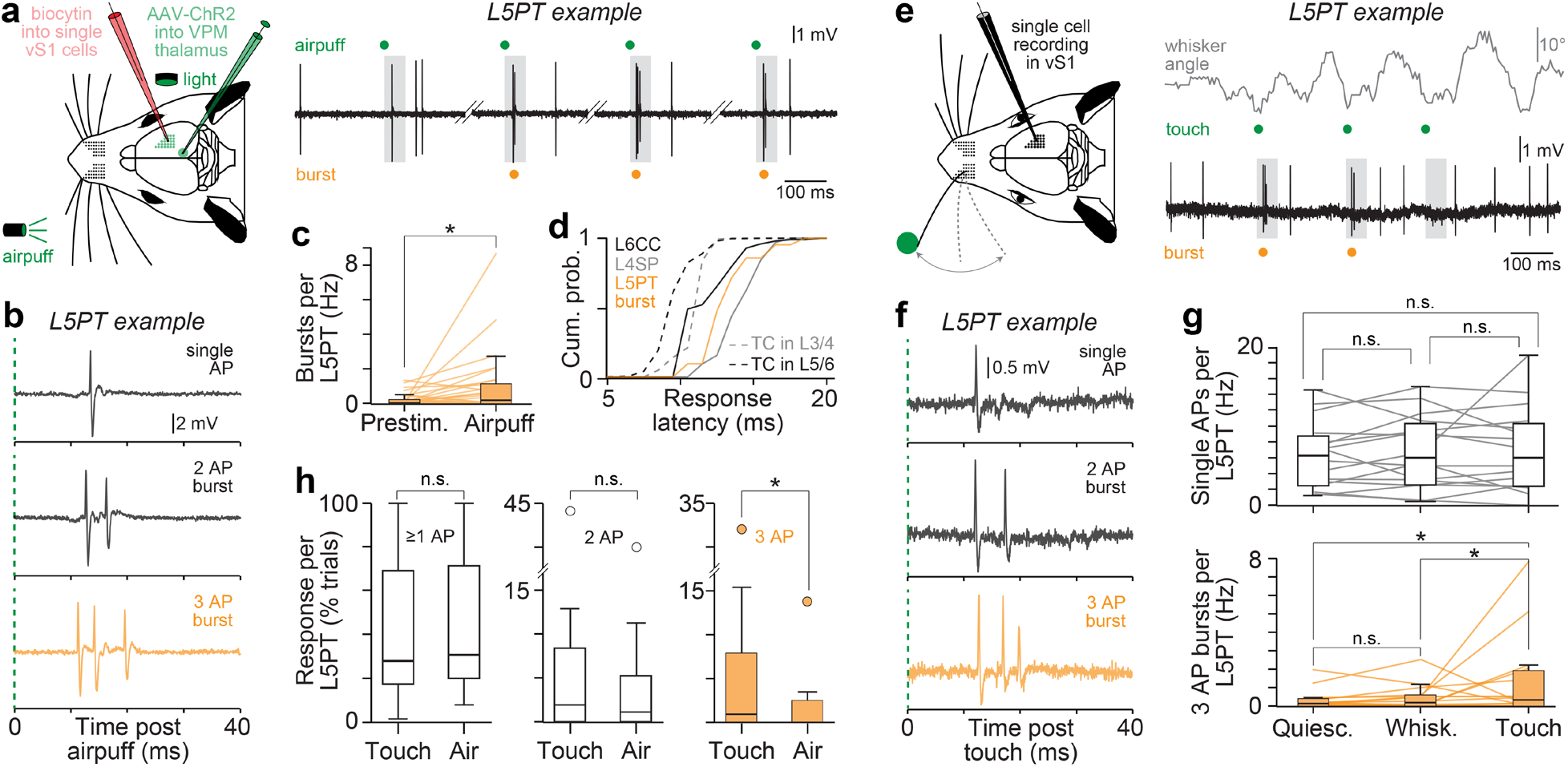
Physiology of TC→L5PT pathway indicates contribution to sensory-evoked bursts. **a**. Left: schematic of the experiments in vS1 of anesthetized rats. Right: example recording for one of the L5PTs in **Fig. 1e** (most right). The trace shows APs recorded during four example trials in which we passively deflected the whiskers by a low-pressure airpuff. The grey boxes denote the first 50 ms after stimulus onset for which we analyzed sensory-evoked responses. **b**. Example trials for the L5PT from panel a show three types of sensory-evoked responses. **c**. Burst rates before (prestim.) and after passive stimuli (n=1,201 trials, N=25 L5PTs). **d**. Latencies of sensory-evoked APs (n=370 trials, N=32 cells) in L6CCs, L4SPs and L5PTs. Latencies of sensory-evoked LFPs show when TC input reaches L5/6 and L3/4 (n=886 trials, N=62 recording depths). **e**. Left: schematic of the experiments in vS1 of awake, head-fixed rats. Right: example recording for one L5PT. The trace shows APs recorded during three example trials in which the animal touched a pole with a single whisker during periods of voluntarily rhythmic whisker movements (whisking). The grey boxes denote the 50 ms time window for which we analyzed sensory-evoked responses. **f**. Example trials of the L5PT from panel e show that active touch evokes the same three types of sensory responses that we observed for passive stimuli in panel b. **g**. Top: single AP rates during periods of quiescence, whisking and active touch (n=1,763 trials, N=16 L5PTs). Bottom: same as in top panel but for bursts with 3APs. **h**. Comparisons of response probabilities (≥1AP), and probabilities for bursts with 2 and 3 APs between passive (airpuff) and active (touch) stimuli.

We tested whether our observations generalize to active sensing conditions. For this purpose, we quantified the activity of L5PTs (n=17) in the barrel cortex of awake rats. Animals were not trained to perform tactile behavior. Instead, sensory responses were evoked by whisker touch of a pole during periods of voluntarily rhythmic whisker movements **(Fig. 2e)**. Active touch stimuli in naïve rats evoked the same three types of responses that we observed in L5PTs for passive stimuli in anesthetized rats **(Fig. 2f)**. Also consistent with the anesthetized data, touch-evoked responses in L5PTs preceded those of L4SPs (Mann-Whitney U: p<0.02). Surprisingly, specifically bursts with 3 APs increased in L5PTs upon touch (ANOVA with multiple comparison: p<0.02), whereas the occurrences of single APs (p>0.65), and of bursts with 2 APs (p>0.08), were not significantly elevated compared to periods preceding the stimulus **(Fig. 2g)**. Moreover, active and passive stimuli evoked virtually identical responses in L5PTs (Mann-Whitney U: p>0.97), but bursts with 3 APs were the exception **(Fig. 2h)**: responses with single APs (p>0.48), and bursts with 2 APs (p>0.45), occurred equally likely during both conditions, whereas burst responses with 3 APs were much more frequent in awake rats (p<0.02). Thus, sensory-evoked bursts with 3 APs may have different origins than those with 2 APs, but their indistinguishably short latencies indicate that direct TC input is likely to contribute to both types of burst responses in L5PTs.

Is TC input necessary, or even sufficient, to drive burst responses in L5PTs? Would this drive rely on wiring specificity in the TC→L5PT pathway? We used multi-scale modelling to address these questions **(Movie S1)**. For this purpose, we recorded responses of 98 neurons in VPM thalamus and of different excitatory (EXC) and inhibitory (INH) cell types across barrel cortex **(Fig. 3a)**. We used these functional data to constrain an anatomically detailed, and empirically validated, network model of barrel cortex (*22*). Embedding of reconstructed L5PTs into the network model thereby provided realistic estimates for which neurons in thalamus and barrel cortex could provide input to these *in vivo* recorded neurons **(Fig. 3b)**, and where along the dendrites these inputs could occur **(Fig. 3c, S2a-c)**. We converted the L5PTs into biophysically-detailed models to capture their empirically observed intrinsic physiology – including calcium APs and burst firing **(Fig. S2d-i)**. Finally, we tuned synaptic strengths to match empirically observed unitary post-synaptic potential (uPSP) distributions **(Fig. S2j)**. We generated 660,960 of such multi-scale models that differed in connectivity patterns, dendrite morphologies, biophysical properties and synaptic strengths to account for parameter degeneracy within and across scales **(Fig. S2a-l)**, while each model met the anatomy and electrophysiology that was observed empirically for L5PTs at subcellular, cellular and network scales **(Methods)**. For each multi-scale model, we simulated how L5PT dendrites transform synaptic inputs evoked by passive whisker stimuli into somatic and dendritic APs. About 10% of the models (n=67,424) predicted responses consistent with those observed *in vivo* **(Fig. S2k)**, including the three response types **(Fig. 3c)**. Even though simulations were not constrained to match this *in vivo* observation, single AP responses occurred also most frequently *in silico*, followed by bursts with 2 APs, whereas bursts with 3 APs were least abundant **(Fig. S2l)**.

**Fig. 3:**
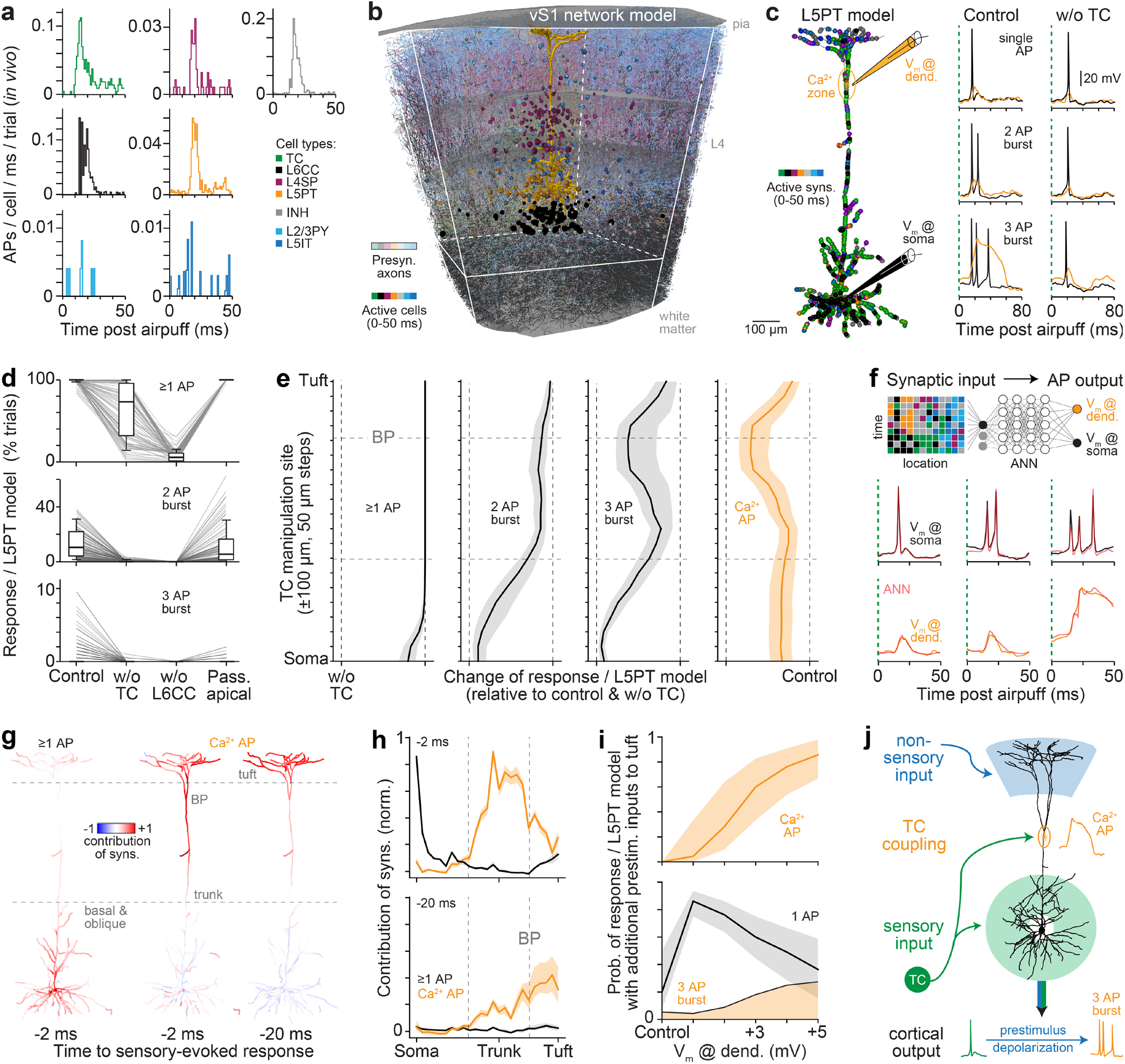
Multi-scale simulations predict TC→L5PT pathway drives sensory-evoked bursts. **a**. Post-stimulus-time-histograms (PSTHs) for passive whisker stimuli for VPM neurons (N=8), pyramidal neurons (PY) in L2/3 (N=8), L4SP (N=10), intratelencephalic neurons (IT) in L5 (N=9), L5PT (N=25), L6CC (N=7), and L2/3-L6 INHs (N=22). **b**. Example of multi-scale model. L5PT whose anatomy and physiology are shown in **Fig. 1e** and **Fig. 2a-b** is converted into biophysically detailed multi-compartmental models **(Fig. S2d-f)** and embedded into network model of vS1. Shaded colors represent axons for one configuration of neurons that are presynaptic to the L5PT. Solid colors represent somata for one configuration of these presynaptic neurons that provide input after stimulus onset. Colors as in panel a. **c**. Left: dendrite distribution of active synapses corresponding to the configuration of active cells in panel b. Right: three example simulation trials for the multi-scale model from panel b with (left) and without TC input (right). The bottom traces represent the configuration of active synapses shown on the left. **d**. *In silico* manipulations (n=16,287,200 trials, N=20,359 models). **e**. *In silico* manipulation in which we removed 50 TC synapses within 100 μm at different dendritic locations, ranging from the soma to the tuft (n=6,340,000 trials, N=317 models). **f**. Top: schematic of ANNs that we trained on the simulations. See **Fig. S3a** for details. Center: Example for how trained ANNs predict membrane potentials (V_m_) at the soma (black) for the three response types (n=8,160 trials, N=408 models). Bottom: same as center panel, but for dendritic calcium domain (orange). **g**. Visualizations for one example multi-scale model show how strongly synapses that are active 2 or 20 ms before somatic (left) or calcium APs (center and right) contribute to these responses. See **Fig. S3b-c** for other time points and morphologies **h**. Synapse contributions depending on soma distance to somatic (black) and dendritic responses (orange). Medians across models (N=68). **i**. *In silico* manipulation: we activated 2 additional synapses per ms along the apical tuft for 20 ms preceding responses (n=4,071,800 trials, N=20,359 models). Additional tuft inputs lead to additional depolarization at the calcium domain at stimulus onset, which increases the probability that L5PTs elicit calcium APs (top). Single AP responses transition into bursts with 3 APs with increasing prestimulus depolarization by tuft inputs (bottom). For this analysis, we included model configurations with bursts of 3 APs in control simulations. **j**. Cartoon of the TC coupling mechanism.

The ability to predict L5PT responses allowed us to explore how the TC→L5PT pathway could contribute. For this purpose, we performed three *in silico* manipulations: we deprived L5PTs from TC input, additionally from L6CC input, or from the active properties of their apical dendrites. Inputs from L6CCs, but not TC input or active apical dendrites, were necessary to evoke L5PT responses **(Fig. 3d)**. These simulations support our previous report that the TC→L6CC→L5PT pathway drives sensory-evoked cortical output via synchronous excitation of proximal dendrites (*14*). However, TC input was necessary to evoke burst responses **(Fig. 3c-d)**. Bursts with 2 APs persisted when apical dendrites were passive, whereas this manipulation abolished bursts with 3 APs. We show why this is the case by locally depriving L5PT models from TC inputs at different dendritic domains **(Fig. 3e)**. TC inputs along proximal dendrites contribute strongly to bursts with 2 and 3 APs. TC inputs to apical trunks – particularly below the primary branch point – contribute strongly to calcium APs, and hence strongly to bursts with 3 APs, but not 2 APs. The simulations support our interpretation of the *in vivo* data **(Fig. 2g-h)** that bursts with 2 and 3 APs have different origins. Thus, all multi-scale models that predicted responses consistent with those observed *in vivo* reached consensus that the TC→L5PT pathway is necessary for sensory-evoked bursts, but that only bursts with 3 APs provide reliable readout for TC-driven calcium APs.

How do other inputs contribute to L5PT responses? How does their interplay with TC inputs cause the transition from single APs to burst responses? To address these questions, we trained artificial neural networks (ANNs; n=68) on the simulations to learn general input-output relationships in L5PTs **(Movie S2)**. The ANNs predicted somatic (median AUROC 0.98) and dendritic (0.96) responses with remarkable precision **(Fig. 3f)**. The ANNs could hence reveal how each synapse – depending on its dendritic location and time of activation – contributes to each response type **(Fig. 3g, S3a)**. Confirming the *in silico* manipulations in **Fig. 3d-e**, synapses along proximal dendrites drive somatic responses, synapses along the apical trunk drive sensory-evoked calcium APs **(Fig. 3h)**. Notably, the contributions that we identified for the time of stimulus onset to generate calcium APs matched the TC→L5PT synapse distribution on apical dendrites **(Fig. 1g)**. The ANNs also showed consistently across all L5PT models with different dendrite morphologies and biophysical properties that apical tuft inputs contribute to calcium APs **(Fig. 3g-h)**, surprisingly for periods up to ∼40 ms preceding stimuli **(Fig. S3b-c)**. We therefore repeated our simulations and activated 2 additional apical tuft synapses per ms for 20 ms preceding responses. This *in silico* manipulation shifted the membrane potential at the calcium domain before response onset closer to threshold by 1-4 mV. This additional prestimulus depolarization by distal tuft inputs increased the probability that TC input drives calcium APs **(Fig. 3i)**, which hence increased the probability that single AP responses transition into bursts with 3 APs – a mechanism which we term ‘TC-coupling’ **(Fig. 3j)**.

How can we test the predicted functions of the TC→L5PT pathway experimentally? The virus we had injected into VPM thalamus also expressed channelrhodopsin (ChR2) in the presynaptic TC terminals, which allowed us to activate TC synapses with 10 ms light pulses while recording *in vivo* from reconstructed and simulated L5PTs **(Fig. 4a)**. Virtually all EXC and INH neurons that we recorded from across barrel cortex (56/61) – including L5PTs (22/24) – showed reliable, fast responses to light stimuli **(Fig. 4b)**. Within the population of L5PTs, light stimuli evoked the same three response types that we had observed for passive and active sensory stimuli. Single APs occurred most frequently (78% of the responses), followed by bursts with 2 APs (21%), which occurred also reliably across the majority of L5PTs (14/22). By contrast, bursts with 3 APs were virtually absent (1%), and occurred only in one of the L5PTs. Thus, supporting the *in silico* predictions **(Fig. 3i)**, TC input is sufficient to drive bursts with 2 APs in L5PTs, but in the absence of additional prestimulus inputs, it is generally insufficient to drive bursts with 3 APs.

**Fig. 4:**
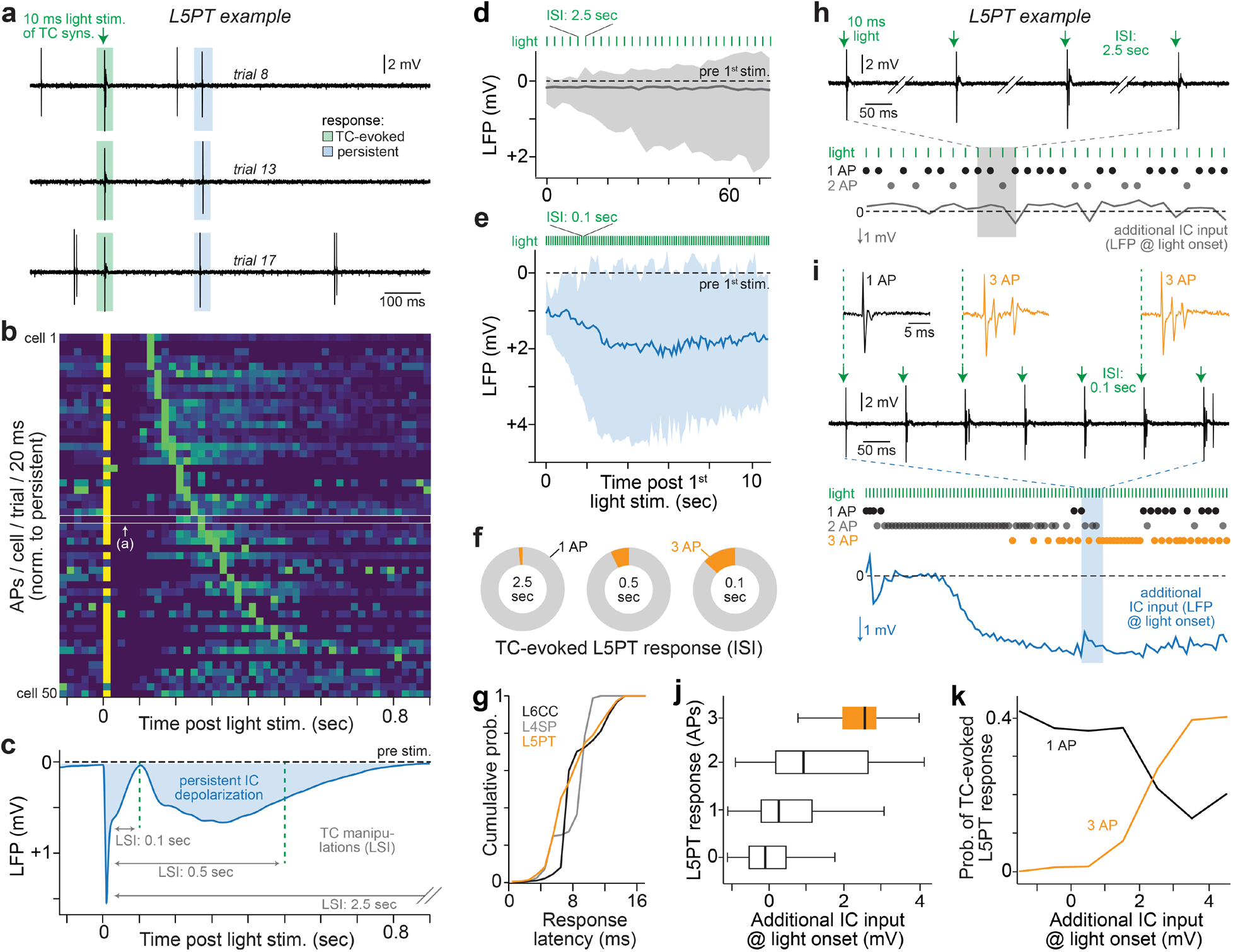
Optogenetic *in vivo* manipulations evoke bursts via the TC→L5PT pathway. **a**. L5PT example whose *in vivo* and *in silico* responses are shown in **Fig. 2a-b** and **Fig. 3c**. The example traces show APs recorded during three trials in which we activated TC synapses by a 10 ms light pulse. **b**. PSTHs for neurons in vS1: L4SP (N=3), L5IT (N=3), L5PT (N=20), L6CC (N=6), L2/3-L6 INH (N=18). Neurons are sorted by persistent activity peak, not by cell type. **c**. Average LFP in vS1 evoked by 10 ms light pulses shows that elevated IC activity from panel b leads to persistent depolarized state across barrel cortex **(Fig. S4a)**. We hence use LFP recordings to estimate the amount of IC inputs that L5PTs receive during *in vivo* manipulations with three LSIs. **d**. LFP at 2.5 sec LSIs (n=894 trails, N=17 rats). **e**. Same as in panel d but for 0.1 sec LSIs indicates that L5PTs receive additional inputs due to elevated IC activity that persists throughout all stimuli. LFP reflects recordings in which we observed bursts with 3 APs (n=750 trials, N=6 L5PTs). **f**. Fraction of light-evoked bursts with 3 APs increases with decreasing LSIs (left to right: 1.2, 5.9, 9.4%). **g**. Latencies of light-evoked responses in L6CCs, L4SPs and L5PTs at 0.1 sec LSIs (n=6,721 trials, N=39 cells). **h**. Example recordings for the L5PT from panel a show four example trials at 2.5 sec LSIs. Bottom panel shows which type of response was elicited by light and the estimated amount of additional IC input (i.e., LFP amplitude within the 10 ms before each light stimulus). **i**. Same L5PT as in panel h, but for 0.1 sec LSIs. Note: bursts with 3 APs did not occur before additional IC input increased. **j**. The TC-evoked response type in L5PTs reflects the additional IC input before a light stimulus (i.e., LFP at 0.1 sec LSIs; n=3,500 trials, N=24 L5PTs). **k**. The higher additional IC input at stimulus onset (n=5,644 trials, N=24 L5PTs), the higher the probability that TC-evoked single APs transition into bursts with 3 APs – supporting the *in silico* prediction in **Fig. 3i**.

Apart from fast responses to light stimuli, most neurons (50 of 61) – independent of their laminar location and cell type – had a second peak of elevated activity, which occurred at later time points that were specific for each neuron **(Fig. 4a-b)**. LFP recordings showed that this wave of elevated activity by IC neurons leads to a depolarized state that persists across all layers of barrel cortex for ∼0.8 seconds **(Fig. 4c, S4a)**. Compared to periods preceding light stimuli, L5PTs hence receive elevated amounts of IC inputs along all dendritic domains, including their tufts **(Fig. S2b-c)** for up to 0.8 seconds. This elevated amount of IC inputs likely provides L5PT dendrites with additional depolarization, which according to our *in silico* predictions for TC-coupling could enable TC input to drive bursts with 3 APs. To test this possibility, we activated TC synapses again (1) when L5PTs receive no additional input **(Fig. 4d)** – i.e., light stimulus intervals (LSIs) are much longer than the duration of elevated IC activity (LSI of 2.5 seconds), (2) when L5PTs receive weak additional input – i.e., light stimuli occur towards the end of elevated IC activity (LSI of 0.5 seconds), or (3) when L5PTs receive strong additional input – i.e., light stimuli occur at the beginning of elevated IC activity (LSI of 0.1 seconds). The two latter *in vivo* manipulations with light stimuli during the depolarized IC state led indeed to TC-driven bursts with 3APs. In particular, LSIs of 0.1 seconds resulted in elevated IC activity – estimated by simultaneously recording LFPs in barrel cortex **(Fig. 4e)** – that persisted throughout all trials. It was in this set of experiments where we observed bursts with 3APs most reliably **(Fig. 4f)**. These L5PT bursts occurred simultaneously with L6CC and L4SP responses (Mann-Whitney U: p>0.4), indicating that they are driven directly by TC input **(Fig. 4g)**. In further support of our *in silico* predictions for TC-coupling, we found that the stronger the additional IC activity was right before a light stimulus – i.e., estimated by LFP amplitude in barrel cortex **(Fig. 4h-i)** – the higher was the probability that a burst with 3 APs occurred **(Fig. 4j)**, which we did not observe in other cell types **(Fig. S4b/c)**. Thus, TC input can drive the transition of L5PT responses from single APs to both types of bursts, but specifically the transition to bursts with 3 APs depends on the amounts of prestimulus inputs by other (here: IC) sources **(Fig. 4k)**.

## Discussion

In this study we sought to reveal why thalamus relays sensory input directly to the major cortical output cells. We found remarkable wiring specificity in the TC→L5PT pathway, and show that it can function to drive fast bursts with 3 APs depending on prestimulus depolarization of the calcium domain **(Fig. 3j)**. Which inputs could provide such depolarization? There is no single answer to this question. Long-range axons from several cortical and subcortical areas innervate barrel cortex layer 1 (*23*), but also diverse local populations from all layers (*24*). Many of these pathways carry nonsensory information (*3*) – e.g. whisker motion or arousal (*25-27*) – and they can contribute to prestimulus depolarization directly or indirectly via disinhibitory circuits (*23, 28, 29*). Moreover, TC-coupling does not rely on precise timing. Instead, any pathway that provides prestimulus inputs – depending on behavioral task and state (*26, 30*) – will contribute to the modulation of sensory-evoked cortical output with bursts. Thus, sensory input drives responses in L5PTs, but nonsensory inputs preceding the stimulus determine which type of response occurs.

This conclusion is supported by our neurophysiological measurements. During anesthesia, where prestimulus inputs by nonsensory pathways are largely absent (*31*), specifically bursts with 3 APs are scarce for both sensory stimuli and activation of TC synapses by light. In turn, despite anesthesia, the occurrence of this response type increases with increasing amounts of prestimulus inputs – here evoked by optogenetic activation of local pathways. Similarly, we show in awake animals that specifically bursts with 3APs increase upon active touch stimuli, which likely reflects prestimulus depolarization by whisker motion, e.g. via inputs from motor cortex that were reported previously to facilitate sensory-evoked calcium APs (*27*). Thus, while it remains an open challenge to simultaneously record from the soma and distal dendrites during behavior, our findings set the stage to dissect how sensory-evoked cortical output encodes nonsensory inputs.

Calcium APs are a general feature of L5PTs across cortex (*3*), and of pyramidal neurons in other brain areas, such as hippocampus (*32*). The current idea for how calcium APs modulate somatic output with bursts was discovered *in vitro*, and is referred to as ‘coincidence detection’ (*5*). In this mechanism, back-propagating APs (bAPs) into the apical trunk depolarize the calcium domain, which opens a short time window for distal inputs to generate bursts. While bursts with 3 APs also represent reliable readout for calcium APs during coincidence detection, our findings indicate that this mechanism is unlikely to account for bursts that occur in L5PTs upon stimulus onset. Although we cannot completely rule out this possibility, the fact that all multi-scale models – which capture the empirically observed features of coincidence detection for a wide range of morphological, biophysical and synaptic properties – reached consensus that bursts with 3APs reflect TC-coupling argues against bAPs and for prestimulus depolarization as their origin. In further support of this conclusion is the fact that bAPs were found to have little influence on apical calcium signals that are recorded *in vivo* (*6, 7, 20, 33*). Instead, these signals are thought to reflect the local activation of calcium channels (*7*), which fits with the wiring specificity we found in the TC→L5PT pathway, and which is the basis for TC-coupling. Thus, complementing coincidence detection, TC-coupling may represent a general circuit theme by which long-range pathways can utilize calcium APs to couple the information they provide with that from anatomically segregated information streams.

Could TC-coupling contribute to the link between perception and sensory-evoked bursts in L5PTs? Output from barrel cortex to higher-order somatosensory thalamus – the posterior medial nucleus (POm) – is critical for perception (*34*). POm thalamus is a major target for L5PTs in barrel cortex (*35*). This corticothalamic pathway forms giant terminals with strong synapses (*36*). Sensory-evoked cortical output thereby reliably activates POm neurons, which then provide feedback to barrel cortex (*17, 29, 37*) and other cortical areas, including frontal cortices (*38*). Due to strong depression of L5PT→POm synapses (*36*), bursts that arrive in isolation are filtered out (*39, 40*). By contrast, when multiple bursts arrive simultaneously, POm neurons encode these bursts in their TC feedback (*41, 42*), which enhances subsequent sensory processing in barrel cortex (*43*), and likely in other cortices that receive POm input (*26*). Here we show that TC-coupling can increase the occurrences of bursts that are simultaneously evoked upon stimulus onset across L5PTs. It is hence tempting to speculate that TC-coupling amplifies the activation of transthalamic pathways, and thereby enhances sensory processing across cortical areas. This theory is consistent with observations in humans, where visual stimuli are only perceived if they evoke responses in primary visual cortex that ‘ignite’ amplified activation across large-scale prefrontal-parietal networks (*44*). Thus, TC-coupling in primary sensory cortices may be first in a cascade of mechanisms by which transthalamic-corticocortical interactions transform sensory input into perception.

## Materials and Methods

### Virus injection

Male Wistar rats were provided by Charles River Laboratories. All experiments were carried out after evaluation by the local German authorities, and in accordance with the animal welfare guidelines of the Max Planck Society. Boutons along primary TC axons were virus labeled as described previously (*14*). Briefly, rats aged 22-25 days (P22-25) were anesthetized with isoflurane supplemented by Caprofen (5mg/ kg) and Buprenorphine SR (1mg/kg) as analgesia, then placed into a stereotaxic frame (Kopf Instruments, model 1900), and provided with a continuous flow of isoflurane/O2 gas. Body temperature was maintained at 37°C by a heating pad. A small craniotomy was made above the left hemisphere 2.85 mm posterior to bregma and 3.2 mm lateral from the midline. The head of the rat was leveled with a precision of 1 μm in both the medial-lateral and anterior-posterior planes using an eLeVeLeR electronic leveling device (Sigmann Electronics, Hüffenhardt, Germany) mounted to an adaptor of the stereotaxic frame. An injecting pipette containing an adeno-associated virus (AAV) was lowered into the VPM thalamus (5.05 mm from the pia). Injection sites were confirmed to be restricted to VPM thalamus by three criteria **(Fig S1)**. The virus (*45*) – rAAV2/1-Syn-hChR2(H134R)-mCherry (titer: 1×10^12^ gc ml^-1^) was provided by Martin Schwarz (University of Bonn, Germany). 50-70 nL of the virus were injected using a 30cc syringe coupled to a calibrated glass injection capillary.

### Electrophysiology in anesthetized animals

All experiments were carried out after evaluation by the local German authorities, and in accordance with the animal welfare guidelines of the Max Planck Society. Non-virus injected rats (P28-48), as well as AAV injected rats (after a 16-21 day incubation period) were anesthetized with urethane (1.8 g/kg body weight) by intraperitoneal injection. The depth of anesthesia was assessed by monitoring pinch withdrawal, eyelid reflexes, and vibrissae movements. Body temperature was maintained at 37°C by a heating pad. Cell-attached recording and biocytin labeling was performed as described in detail previously (*46*). Briefly, a small craniotomy was made above the left hemisphere 2.5 mm posterior and 5.5 mm lateral to the bregma (for recordings in vS1), or 2.9-3.5 mm posterior to the bregma, 2.4-3.4 mm lateral from the midline, and at 5-6 mm depth from the pia for recordings in VPM thalamus. APs were recorded using an extracellular loose patch amplifier (ELC-01X, npi electronic GmbH) or an Axoclamp 2B amplifier (Axon instruments, Union City, CA, USA), digitized using a CED power1401 data acquisition board (CED, Cambridge Electronic Design, Cambridge, UK), and low pass filtered (300 Hz) to measure the LFP. APs and LFPs were recorded before and during 20-30 trials of caudal multi-whisker deflections by a 700 ms airpuff (10 PSI), delivered through a 1 mm plastic tube from a distance of 8-10 cm from the whisker pad (*35*). Stimulation was repeated at 2.5 sec intervals. We manually validated each detected AP. We assigned trials to the response types depending on their activity within the first 50 ms after stimulus onset: 3 AP bursts were defined as 3 APs occurring within 30 ms, 2 AP bursts as 2 APs within 10 ms. Trials that fulfilled both criteria were assigned as 3 AP bursts. The same criteria were used for classifying optogenetic responses, active touch responses, and the simulations. We determined the latencies of TC onset at different cortical depths in vS1 by detecting the first depolarization of the LFP after stimulation. In AAV injected rats, optical stimulation of ChR2-expressing TC terminals was provided by a 400 μm diameter optical fiber (ThorLabs #RJPSF2) coupled to a 470 nm wavelength LED source (ThorLabs M470F3) and powered by an LED driver (ThorLabs #DC2200). A 10 ms pulse of light generated 1 mW output power at the end of the optical fiber as measured by a laser power meter (ThorLabs #PM100A) coupled to a photodiode (ThorLabs #S121C). The optical fiber was positioned using a 3-axis motorized micromanipulator (Luigs and Neuman) approximately 1-2 mm above the cortical surface, so that the light beam resulted in a 1-2 mm disc of light above the recording site in vS1. Control of the LED driver was implemented with Spike2 software (CED, Cambridge, UK.). APs and LFPs were recorded during 20-100 trials of 10 ms light pulses, at light stimulus intervals of 2.5, 0.5 and 0.1 seconds. Following the electrophysiological measurements, neurons were labeled with biocytin. Labeling sessions were repeated several times. After 1-2 hours for biocytin diffusion, animals were transcardially perfused with 0.1M phosphate buffer (PB) followed by 4% paraformaldehyde (PFA). Brains were removed and post-fixed with 4% PFA for 8 to 12 hours, transferred to 0.1 M PB and stored at 4°C.

### Electrophysiology in awake animals

All experiments were carried out in accordance with the animal welfare guidelines of the VU Amsterdam, the Netherlands, as described previously (*47*). Briefly, male Wistar rats (P39 ± 4, provided by Charles River Laboratories) were positioned in the recording setup using a head-post. During surgical preparation, rats were anesthetized using 1.6% isoflurane in 0.4 l/h O2 + 0.7 l/h NO2 and depth of anesthesia was assured by the absence of foot and eyelid reflexes. In addition, post-operative analgesia (buprenorphine, 0.1–0.5 mg/kg) was given. Body temperature was maintained at 37 °C with a heating pad. In the week prior to surgery, rats were handled daily to accustom them to the experimenter and housed in pairs in enriched cages. In the week after surgical preparation, rats were head-fixed twice per day for 2–3 days in preparation of the recording session. Rats quickly adjusted to the head-fixation period, allowing stable recording configurations without the need for body restraint. On the recording day, rats were anaesthetized with isoflurane (1.25% in 0.4 l/h O2 + 0.7 l/h NO2), and targeted loose-patch recordings were made using intrinsic optical imaging. Passive whisker stimulation and receptive field mapping were used to confirm intrinsic optical imaging results. Afterwards, whiskers were clipped to 5 mm, except the principal or a single surround whisker and anesthesia was terminated. Rats woke up from isoflurane anesthesia within several minutes and APs during active object touch were quantified only after rats were fully awake, monitored by body posture and exploratory whisking. Active touch resulted from whisker self-motion and was monitored with high-speed videography (375 seconds continuously at 200 frames/sec, MotionScope M3 camera, IDT Europe, Belgium). The object was positioned 2 cm lateral from the whisker pad and anterior relative to the whisker set point (obtained during quiescent episodes). This ensured that touches were the consequence of whisker protraction. Touch events were detected manually frame-by-frame. After the electrophysiological measurements, neurons were labeled with biocytin as described above, animals were deeply anaesthetized with urethane (>2.0 g/kg) and perfused with 0.9% NaCl, then 4% paraformaldehyde (PFA). Brains were post-fixed in 4% PFA overnight at 4 °C and transferred to 0.9% NaCl.

### Histology

For recordings in vS1 of non-virus injected and awake rats, 22-25 consecutive 100 μm thick vibratome sections were cut tangentially to vS1 (45° angle) ranging from the pial surface to the white matter, or coronally (for recordings in VPM thalamus). All sections were treated with avidin-biotin (ABC) solution, and subsequently neurons were identified using the chromogen 3,3′-diaminobenzidine tetrahydrochloride (DAB). All sections were mounted on glass slides, embedded with Mowiol, and enclosed with a cover slip. For recordings in AAV injected rats, the recorded hemisphere was cut tangentially into 45-48 consecutive 50 μm thick vibratome sections from the pial surface to the white matter. The remaining brain tissue was embedded in 10% gelatin (Sigma Aldrich #G2500) and cut coronally into consecutive 100 μm thick sections to identify the virus injection site. Sections were first washed three times in 0.1 M PB and treated with streptavidin conjugated to AlexaFluor488 (5μg/ml) (Molecular Probes #S11223) in 0.1 M PB containing 0.3% Triton X-100 (TX) (Sigma Aldrich #9002-93-1), 400 μl per section for 3-5 hours at room temperature in order to visualize biocytin labeled neuronal structures. To enhance the fluorescence expressed by the virus and to label primary thalamic synapses, slices were then double immunolabeled with anti-mCherry antibody and anti-VGlut2 antibody. Sections were permeabilized and blocked in 0.5% Triton x-100 (TX) (Sigma Aldrich #9002-93-1) in 100 mM PB containing 4% normal goat serum (NGS) (Jackson ImmunoResearch Laboratories #005-000-121) for 2 hours at room temperature. The primary antibodies were diluted 1:500 (Rabbit anti-mCherry, Invitrogen #PA5-34974 Invitrogen #M11217 and mouse anti-VGlut2 antibody, Synaptic Systems #135421) in PB containing 1% NGS for 48 hours at 4°C. The secondary antibodies were diluted 1:500 (goat anti-Rabbit IgG Alexa-647 H+L Invitrogen #A21245 and goat anti-Mouse IgG Alexa-405 H+L Invitrogen #A31553), and were incubated for 2-3 hours at room temperature in PB containing 3% NGS and 0.3% TX. All sections were mounted on glass slides, embedded with SlowFade Gold (Invitrogen #S36936) and enclosed with a coverslip.

### Morphological reconstructions

Neuronal structures were extracted from image stacks using a previously reported automated tracing software (*48*). For reconstruction of fluorescently labeled neurons and locating of AAV labeled TC synapses, images were acquired using a confocal laser scanning system (SP5; Leica Microsystems). 3D image stacks of up to 2.5 mm × 2.5 mm × 0.05 mm were acquired at 0.092 × 0.092 × 0.5 μm per voxel (63x magnification, NA 1.3). Image stacks were acquired for each of 45-48 consecutive 50 μm thick tangential brain slices that range from the pial surface to the white matter. Manual proof-editing of individual sections, and automated alignment across sections were performed using custom-designed software (*49*). Pia, barrel and white matter outlines were manually drawn on low-resolution images (4x magnification dry objective). Using these anatomical reference structures, all reconstructed morphologies were registered to a standardized 3D reference frame of rat vS1 (*50*). The distance from the pial surface to the soma, and 20 morphological features that have previously been shown to separate between excitatory cell types in rat vS1 (*51*) were calculated for each registered morphology **(Fig. S1c)**. For identification of putative TC synapses, biocytin labeled morphologies and AAV labeled VPM terminals were imaged simultaneously using the confocal system as described above: biocytin Alexa-488 (excited at 488 nm, emission detection range 495-550 nm), AAV Alexa-647 (excited at 633 nm, emission detection range 650-785 nm). These dual-channel image stacks were loaded into Amira software (Thermo Scientific) for visualization. All reconstructed dendrites were manually inspected, and landmarks were placed on each spine head if a spine head was overlapping with a VPM bouton, to mark the location of a putative synapse. The shortest distance of each landmark to the dendrite reconstruction was determined, and the path length distance was calculated from that location along the reconstructed neuron to the soma. For validation of putative TC synapses, image stacks were acquired via super-resolution microscopy (SP8 LIGHTNING; Leica Microsystems) via a glycerol/oil immersion objective (HCX PL APO 63x magnification, NA 1.3), a tandem scanning system (8 kHz resonance scanning speed), and spectral detectors with hybrid technology (GaAsP photocathode; 8x line average): VGlut2 Alexa-405 (excited at 405 nm, emission detection range: 410-480 nm), biocytin Alexa-488 (excited at 488 nm, emission detection range 495-550 nm), AAV Alexa-647 (excited at 633 nm, emission detection range 650-785 nm). Triple-channel image stacks were acquired at 29.5 × 29.5 × 130 nm per voxel. Image stacks were visualized in Amira, and manually inspected for VGlut2 at contact sites between spines and AAV-infected TC boutons within single optical sections (range of VGlut2-positive contacts: 68-100%, n=349 contacts across 26 basal and apical trunk dendrites of 8 L5PTs).

### Multi-compartmental models

We selected three morphologies from the *in vivo* recorded L5PTs with reconstructed TC synapses that capture the range of morphological variability (i.e., the L5PTs with the deepest and most superficial BP, and one L5PT from the bulk of the distribution). Multi-compartmental models were generated for these dendrite morphologies as described previously (13, 36). Briefly, a simplified axon morphology was attached to the soma of the reconstructed L5PT morphology (44). The axon consisted of an axon hillock with a diameter tapering from 3 μm to 1.75 μm over a length of 20 μm, an axon initial segment of 30 μm length and diameter tapering from 1.75 μm to 1 μm diameter, and 1 mm of myelinated axon (diameter of 1 μm). Next, a multi-objective evolutionary algorithm was used to find parameters for the passive leak conductance and the density of Hodgkin-Huxley type ion channels on soma, basal dendrite, apical dendrite and axon initial segment, such that the neuron model is able to reproduce characteristic electrophysiological responses to somatic and dendritic current injections of L5PTs within the experimentally observed variability, including bAPs, calcium APs, and AP responses to prolonged somatic current injections (36). We augmented the original biophysical model of L5PTs (13, 36) with two ion channel parameters: according to a previous report (45), the density of the fast non-inactivating potassium channels (Kv3.1) was allowed to linearly decrease with soma distance until it reaches a minimum density (i.e., the slope and minimum density are two additional parameters). The diameter of the apical dendrites was optimized by a scaling factor between 0.3 and 3. We incorporated the IBEA algorithm (46) for optimization. The optimization was terminated if there was no progress or when acceptable models had been found. We repeated the optimization process several times and generated in total in 909722, 292151 and 201028 acceptable models for the L5PT with the most superficial, deepest and in-between BP locations, respectively. Each of these models reproduce the characteristic electrophysiology of L5PTs, but utilized largely different ion channel distribution to do so **(Fig. S2d-i)**. A subset of models were additionally constrained to be able to perform coincidence detection multiple times within 200 ms. From each optimization run, we selected one model for which the maximal deviation from the mean of the empirical data in units of standard deviation across all objectives was minimal (0.9-1.9 mean STDs across objectives). Additionally, from the whole data base of models, we selected models that are most diverse in the active ion currents contributing to their calcium APs for each morphology. To do so, we quantified the charge exchanged through active conductances for each model during the calcium AP and computed the contributions of different ion channels to the total hyperpolarizing or depolarizing currents. For each ion channel and each morphology, we selected the model in which the contribution of the respective channel was largest, which resulted in a set of models utilizing completely disparate mechanisms to generate dendritic calcium APs **(Fig. S2d-h)**. We selected 68 from these models for network embedding and simulations as described below (most superficial BP: 40, in-between BP: 13, deepest BP: 15).

### Network models

TC synapse positions along the L5PT models were measured empirically as described above. Cell type-specific IC excitatory connections are derived from an anatomically realistic network model of rat vS1 (*22*), a procedure, which has been described in detail previously (*14*). We registered the dendrite morphologies selected for multi-compartmental modelling in the network model at nine locations within the cortical barrel column representing the C2 whisker, which is located approximately in the center of vS1, while preserving its (*in vivo*) soma depth (*50*). The locations were the column center and equally spaced angular intervals with a distance of ∼100 μm to the column center. For each of the nine locations, we estimated the numbers and dendritic locations of cell type-specific synapses that impinge onto the dendrites of the L5PT model. Finally, IC inhibitory synapses were distributed randomly on the dendritic tree. Synapses were assigned to presynaptic neurons in the following manner. TC synapses: we randomly assigned each TC synapse to 350 TC neurons, which is the average number of neurons in the somatotopically aligned C2 barreloid (*52*). IC excitatory synapses: presynaptic neuron IDs are an output of the network model (*22*). IC inhibitory synapses: each synapse was assigned to one virtual neuron.

### Synapse models

Synapse models and synaptic parameters (rise and decay times, release probabilities, reversal potentials) were reported previously (*14*). Briefly, conductance-based synapses were modeled with a double-exponential time course. Excitatory synapses contained both AMPA receptors (AMPARs) and NMDARs. Inhibitory synapses contained GABA_A_Rs. The peak conductance of excitatory synapses from different presynaptic cell types was determined by assigning the same peak conductance to all synapses of the same cell type, activating all connections of the same cell type (i.e., all synapses originating from the same presynaptic neurons) one at a time, and comparing parameters of the resulting unitary postsynaptic potential (uPSP) amplitude distribution (mean, median and maximum) for a fixed peak conductance with experimental measurements *in vitro* (IC input (*53*)) or *in vivo* (TC input (*12*), **Fig. S2j**). The peak conductance for synaptic inputs from each cell type was systematically varied until the squared differences between parameters of the *in silico* and *in vitro/in vivo* uPSP amplitude distributions were minimized (i.e., the mean, median and maximum of the distributions were used, and mean and median were weighted twice relative to the maximum). This procedure was repeated for each multi-compartmental model using the connectivity model for the location in the center of the C2 column. The peak conductance at inhibitory synapses was fixed at 1 nS (*54*).

### Simulations

Simulations were performed using Python 3.8, dask (*55*) and NEURON 7.8 (*56*). First, we determined model configurations that result in simulations with AP rates that are within the range observed across L5PTs during periods preceding passive whisker stimuli. For the 612 network-embedded L5PT models (i.e., 68 multi-compartmental models for 3 morphologies and 9 embeddings into the network model, respectively) we simulated prestimulus activity by activating neurons in the network model that are presynaptic to the L5PTs with the spontaneous firing rates that we recorded in anesthetized animals across layers and for different excitatory and inhibitory cell types, and in VPM thalamus **(Fig. 3a)**. We distributed 5000 inhibitory synapses uniformly across the dendritic tree, and activated them with different multiples of the *in vivo* observed firing rates (i.e., 0.25 to 3.75 times the *in vivo* rates, in 0.25 steps). For each of these 9180 model configurations (i.e., 68 L5PT models * 9 network embeddings * 15 EXC/INH ratios), we simulated 48 trials of 1245 ms duration, of which we discarded the first 245 ms as initialization of the simulations. 2241 of the 9180 configurations predicted prestimulus AP rates as observed for L5PTs *in vivo* (i.e., >0 Hz and ≤11 Hz). Second, we determined which of these 2241 model configurations result also in simulations with single AP, 2 AP and 3 AP burst rates that are within the respective ranges observed across L5PTs during the onset response evoked by passive whisker stimuli. For this purpose, we simulated 445 ms of prestimulus activity as described above, and then activated neurons in the network model that are presynaptic to the L5PT models by generating Poisson spike trains for each presynaptic neuron based on the empirically measured PSTHs of the respective cell types **(Fig. 3a)**. We systematically tested whether our simulation results are robust with respect to different stimuli, different timings and ratios of sensory-evoked inhibition relative to excitation. Stimuli: principal whisker, three whiskers within the same row, or three whiskers within two adjacent rows, respectively. Timings: latencies as observed *in vivo*, inhibition shifted by 1 or 2 ms to earlier time points. Ratios: we activated inhibitory neurons with different multiples of the *in vivo* observed firing rates (i.e., 0.1 to 1.0 times the *in vivo* PSTHs, in 0.1 steps). For each of these 161352 model configurations (i.e., 2241 models with realistic prestimulus rates * 3 stimuli * 3 EXC/INH timings * 8 EXC/INH ratios), we simulated 20 trials to select all configurations that predict sensory-evoked responses within the *in vivo* observed ranges (i.e., ≥1 AP probability: >0%, 2 AP burst probability >0 % and ≤40%, 3 AP burst probability ≥0 % and ≤25%). 67424 of the 161352 configurations predicted pre- and post-stimulus activity as we had observed for L5PTs *in vivo*, out of which 22850 configurations contained sensory-evoked bursts. Third, we selected these 22850 model configurations and repeated the simulations, this time 200 instead of 20 trials per model configuration. Moreover, in the first coarse selection step, we had accepted 3 AP burst probabilities up to 25%. Now, we only selected those model configurations with 3 AP burst probabilities up to 13%, which is the maximal value observed in anesthetized animals. The simulations hence identified 20359 model configurations that predict pre- and post-stimulus activity, including fast sensory-evoked bursts with 2 and 3 APs, consistent with the *in vivo* data. These models comprised configurations for all three morphologies (most superficial BP: 12589, in-between BP: 3298, deepest BP: 4472 models), all nine network locations, all 68 biophysical parameter sets, all 3 whisker stimuli, all inhibitory timings, 7/8 post-stimulus EXC/INH ratios, and 13/15 pre-stimulus EXC/INH ratios. Finally, we used these 20359 model configurations for *in silico* manipulations and for training ANNs. For this purpose, we replayed the simulations multiple times, but (1) inactivated sensory-evoked TC input, (2) inactivated additionally L6CC input, (3) removed active conductances along the apical dendrite, (4) or activated 50 additional tuft synapses randomly within 20 ms preceding responses. To investigate distance-dependent impacts of TC synapse distributions, we selected models with 3 AP burst probability >= 2%, and simulated 1000 trials for each of them. These models comprised configurations for all three morphologies (most superficial BP: 280, in-between BP: 1, deepest BP: 36 models). We replayed the simulations 20 times, each time removing 50 TC synapse activations in 200 μm intervals with increasing soma distance ranging from 0-200 to 950-1150 μm.

### ANN training

We used pytorch and python 3.8 to train ANNs on the 200 trials that we simulated for each of the 67424 configurations that predicted pre- and post-stimulus activity as observed for L5PTs *in vivo* (i.e., including the 22850 configurations with sensory-evoked bursts). ANNs were trained to predict the membrane potential at the soma or BP for 60 ms post-stimulus with 1 ms resolution. Training was done in a supervised fashion based on the 200 * 67424 simulations, separately for each of the 68 multi-compartmental L5PT models. Input to the ANNs contained the spatiotemporal synaptic input pattern of 80 ms preceding the time point for which the ANNs should predict the membrane potentials, and the time to the previous somatic and dendritic AP. Output was the membrane potential at the BP. The ANN was applied convolutionally over the 60 ms time period, during which gradients were accumulated. We used a mean-squared-error loss to compare predicted with the simulated membrane potential. Spatiotemporal input patterns were generated by binning synapse activations for each simulation trial based on time and dendritic location for EXC and INH synapses, respectively. Spatial bins were generated by uniformly splitting all dendritic segments (i.e., stretches between branch points) to achieve a bin size closest to 50 μm (e.g. a 90 μm segment is split into two 45 μm bins). Temporal bin size was 1 ms. The ANN is based on a multi-layer-perceptron architecture **(Fig. S3a)**. Layer 1 is linear with input dimensions matching the dimension of the spatiotemporal input patterns: 80 temporal bins * 238, 338 or 365 spatial bins (most superficial, in-between or deepest BP) * 2 synapse types (EXC and INH). Its output dimension is one (i.e., ‘bottleneck’). This dimensionality reduction of the spatiotemporal input patterns to one was sufficient to learn somatic and dendritic responses, which only marginally improved by adding more nodes to the bottleneck (data for 2-4 nodes not show). We concatenated the time to the previous somatic and dendritic AP to the bottleneck. This allowed ANNs to learn intracellular dynamics (e.g. refractory periods), which resulted in higher prediction accuracy. Behind the bottleneck, the ANNs have five ReLU layers with width 40, the last of them provides input to a linear output layer with output dimension 1. All weights in the ANNs were subjected to a L2 regularization. Additionally, we introduced a total variation loss in layer 1, which introduces a bias that favors similar weights for adjacent time points. We used the ‘ADAM’ optimizer for training while decreasing learning rates to assure convergence. For each of the 68 multi-compartmental L5PT models, the simulations were split in a training and test dataset, the latter containing 20000 trials. We tracked training and test loss, and the Pearson correlation coefficient for the test dataset during training. We selected the model with the highest Pearson correlation coefficient. Weights between the input layer and bottleneck reflect the influence of EXC and INH synapses by location and time. We visualized these weights by color-coding the dendrite according to the weights assigned to its spatial and temporal bins **(Fig. S3b-c)**. The model and simulation routines, a documentation of all parameters and the analysis routines are available at ModelDB – https://senselab.med.yale.edu/ModelDB/; accession number: 267166).

### Quantification and statistical analysis

All data are reported as mean ± standard deviation (STD) unless mentioned otherwise. All of the statistical details can be found in the figure legends, figures, and Results, including the statistical tests used, sample size n, and what the sample size represents (e.g. number of animals, number of cells). Significance was defined for p-values smaller than 0.05. Boxplots show median, 25% and 75% percentile; whiskers extend up to 1.5 times the interquartile range. All data beyond the whiskers are shown as outliers. All tests were performed using the scipy python package (version 0.18.2).

## Acknowledgments

We thank Bert Sakmann, Peter Strick and Idan Segev for comments, Martin Schwarz for providing the virus construct and Philipp Harth for visualization of the cortex model.

## Author contributions

MO conceived and designed the study. JMG performed the *in vivo* experiments and discovered the wiring specificity. AB performed the simulations and discovered TC-coupling mechanism. JMG, AB, RF and MO analyzed the data. RN and CPJdK contributed data. MO wrote the paper with AB and JMG.

## References

1. J. Aru, M. Suzuki, M. E. Larkum, Cellular Mechanisms of Conscious Processing. Trends Cogn Sci 24, 814–825 (2020).

2. M. Larkum, A cellular mechanism for cortical associations: an organizing principle for the cerebral cortex. Trends Neurosci 36, 141–151 (2013).

3. K. D. Harris, G. M. Shepherd, The neocortical circuit: themes and variations. Nat Neurosci 18, 170–181 (2015).

4. J. Schiller, Y. Schiller, G. Stuart, B. Sakmann, Calcium action potentials restricted to distal apical dendrites of rat neocortical pyramidal neurons. J Physiol 505 (Pt 3), 605–616 (1997).

5. M. E. Larkum, J. J. Zhu, B. Sakmann, A new cellular mechanism for coupling inputs arriving at different cortical layers. Nature 398, 338–341 (1999).

6. N. Takahashi et al., Active dendritic currents gate descending cortical outputs in perception. Nat Neurosci 23, 1277–1285 (2020).

7. N. Takahashi, T. G. Oertner, P. Hegemann, M. E. Larkum, Active cortical dendrites modulate perception. Science 354, 1587–1590 (2016).

8. R. J. Douglas, K. A. Martin, Neuronal circuits of the neocortex. Annu Rev Neurosci 27, 419–451 (2004).

9. R. J. Douglas, K. A. Martin, in Handbook of Brain Microcircuits, G. M. Shepherd, S. Grillner, Eds. (Oxford University Press, 2010).

10. C. D. Gilbert, T. N. Wiesel, Morphology and intracortical projections of functionally characterised neurones in the cat visual cortex. Nature 280, 120–125 (1979).

11. M. Armstrong-James, K. Fox, Spatiotemporal convergence and divergence in the rat S1 “barrel” cortex. J Comp Neurol 263, 265–281 (1987).

12. C. M. Constantinople, R. M. Bruno, Deep cortical layers are activated directly by thalamus. Science 340, 1591–1594 (2013).

13. C. P. de Kock, R. M. Bruno, H. Spors, B. Sakmann, Layer- and cell-type-specific suprathreshold stimulus representation in rat primary somatosensory cortex. J Physiol 581, 139–154 (2007).

14. R. Egger et al., Cortical Output Is Gated by Horizontally Projecting Neurons in the Deep Layers. Neuron 105, 122–137 e128 (2020).

15. M. Ito, Simultaneous visualization of cortical barrels and horseradish peroxidase-injected layer 5b vibrissa neurones in the rat. J Physiol 454, 247–265 (1992).

16. R. M. Bruno, B. Sakmann, Cortex is driven by weak but synchronously active thalamocortical synapses. Science 312, 1622–1627 (2006).

17. L. Petreanu, T. Mao, S. M. Sternson, K. Svoboda, The subcellular organization of neocortical excitatory connections. Nature 457, 1142–1145 (2009).

18. J. C. Rah et al., Thalamocortical input onto layer 5 pyramidal neurons measured using quantitative large-scale array tomography. Front Neural Circuits 7, 177 (2013).

19. R. Yuste, M. J. Gutnick, D. Saar, K. R. Delaney, D. W. Tank, Ca2+ accumulations in dendrites of neocortical pyramidal neurons: an apical band and evidence for two functional compartments. Neuron 13, 23–43 (1994).

20. F. Helmchen, K. Svoboda, W. Denk, D. W. Tank, In vivo dendritic calcium dynamics in deep-layer cortical pyramidal neurons. Nat Neurosci 2, 989–996 (1999).

21. H. G. Kim, B. W. Connors, Apical dendrites of the neocortex: correlation between sodium- and calcium-dependent spiking and pyramidal cell morphology. J Neurosci 13, 5301–5311 (1993).

22. D. Udvary et al., The impact of neuron morphology on cortical network architecture. Cell Rep 39, 110677 (2022).

23. J. M. Ledderose et al., Layer 1 of somatosensory cortex: An important site for input to a tiny cortical compartment. bioRxiv, 2021.2011.2026.469979 (2023).

24. R. T. Narayanan et al., Beyond Columnar Organization: Cell Type- and Target Layer-Specific Principles of Horizontal Axon Projection Patterns in Rat Vibrissal Cortex. Cereb Cortex 25, 4450–4468 (2015).

25. M. Oberlaender et al., Three-dimensional axon morphologies of individual layer 5 neurons indicate cell type-specific intracortical pathways for whisker motion and touch. Proc Natl Acad Sci U S A 108, 4188–4193 (2011).

26. G. H. Petty, A. K. Kinnischtzke, Y. K. Hong, R. M. Bruno, Effects of arousal and movement on secondary somatosensory and visual thalamus. Elife 10, (2021).

27. N. L. Xu et al., Nonlinear dendritic integration of sensory and motor input during an active sensing task. Nature 492, 247–251 (2012).

28. N. J. Audette, S. M. Bernhard, A. Ray, L. T. Stewart, A. L. Barth, Rapid Plasticity of Higher-Order Thalamocortical Inputs during Sensory Learning. Neuron 103, 277–291 e274 (2019).

29. L. E. Williams, A. Holtmaat, Higher-Order Thalamocortical Inputs Gate Synaptic Long-Term Potentiation via Disinhibition. Neuron 101, 91–102 e104 (2019).

30. J. L. Chen, S. Carta, J. Soldado-Magraner, B. L. Schneider, F. Helmchen, Behaviour-dependent recruitment of long-range projection neurons in somatosensory cortex. Nature 499, 336–340 (2013).

31. M. Suzuki, M. E. Larkum, General Anesthesia Decouples Cortical Pyramidal Neurons. Cell 180, 666–676 e613 (2020).

32. J. Magee, D. Hoffman, C. Colbert, D. Johnston, Electrical and calcium signaling in dendrites of hippocampal pyramidal neurons. Annu Rev Physiol 60, 327–346 (1998).

33. M. Murayama, E. Perez-Garci, H. R. Luscher, M. E. Larkum, Fiberoptic system for recording dendritic calcium signals in layer 5 neocortical pyramidal cells in freely moving rats. J Neurophysiol 98, 1791–1805 (2007).

34. J. Qi, C. Ye, S. Naskar, A. R. Inacio, S. Lee, Posteromedial thalamic nucleus activity significantly contributes to perceptual discrimination. PLoS Biol 20, e3001896 (2022).

35. G. Rojas-Piloni et al., Relationships between structure, in vivo function and long-range axonal target of cortical pyramidal tract neurons. Nat Commun 8, 870 (2017).

36. A. Groh, C. P. de Kock, V. C. Wimmer, B. Sakmann, T. Kuner, Driver or coincidence detector: modal switch of a corticothalamic giant synapse controlled by spontaneous activity and short-term depression. J Neurosci 28, 9652–9663 (2008).

37. V. C. Wimmer, R. M. Bruno, C. P. de Kock, T. Kuner, B. Sakmann, Dimensions of a projection column and architecture of VPM and POm axons in rat vibrissal cortex. Cereb Cortex 20, 2265–2276 (2010).

38. M. Deschenes, P. Veinante, Z. W. Zhang, The organization of corticothalamic projections: reciprocity versus parity. Brain Res Brain Res Rev 28, 286–308 (1998).

39. R. A. Mease, P. Krieger, A. Groh, Cortical control of adaptation and sensory relay mode in the thalamus. Proc Natl Acad Sci U S A 111, 6798–6803 (2014).

40. R. A. Mease, A. Sumser, B. Sakmann, A. Groh, Corticothalamic Spike Transfer via the L5B-POm Pathway in vivo. Cereb Cortex 26, 3461–3475 (2016).

41. R. A. Mease, T. Kuner, A. L. Fairhall, A. Groh, Multiplexed Spike Coding and Adaptation in the Thalamus. Cell Rep 19, 1130–1140 (2017).

42. R. A. Mease, A. Sumser, B. Sakmann, A. Groh, Cortical Dependence of Whisker Responses in Posterior Medial Thalamus In Vivo. Cereb Cortex 26, 3534–3543 (2016).

43. R. A. Mease, M. Metz, A. Groh, Cortical Sensory Responses Are Enhanced by the Higher-Order Thalamus. Cell Rep 14, 208–215 (2016).

44. S. Dehaene, J. P. Changeux, Experimental and theoretical approaches to conscious processing. Neuron 70, 200–227 (2011).

45. F. J. Meye et al., Shifted pallidal co-release of GABA and glutamate in habenula drives cocaine withdrawal and relapse. Nat Neurosci 19, 1019–1024 (2016).

46. R. T. Narayanan et al., Juxtasomal biocytin labeling to study the structure-function relationship of individual cortical neurons. J Vis Exp, e51359 (2014).

47. C. P. De Kock et al., High-frequency burst spiking in layer 5 thick-tufted pyramids of rat primary somatosensory cortex encodes exploratory touch. Communications Biology, (2021).

48. M. Oberlaender, R. M. Bruno, B. Sakmann, P. J. Broser, Transmitted light brightfield mosaic microscopy for three-dimensional tracing of single neuron morphology. J Biomed Opt 12, 064029 (2007).

49. V. J. Dercksen, H. C. Hege, M. Oberlaender, The Filament Editor: an interactive software environment for visualization, proof-editing and analysis of 3D neuron morphology. Neuroinformatics 12, 325–339 (2014).

50. R. Egger, R. T. Narayanan, M. Helmstaedter, C. P. de Kock, M. Oberlaender, 3D reconstruction and standardization of the rat vibrissal cortex for precise registration of single neuron morphology. PLoS Comput Biol 8, e1002837 (2012).

51. M. Oberlaender et al., Cell type-specific three-dimensional structure of thalamocortical circuits in a column of rat vibrissal cortex. Cereb Cortex 22, 2375–2391 (2012).

52. H. S. Meyer et al., Cellular organization of cortical barrel columns is whisker-specific. Proc Natl Acad Sci U S A 110, 19113–19118 (2013).

53. P. Schnepel, A. Kumar, M. Zohar, A. Aertsen, C. Boucsein, Physiology and Impact of Horizontal Connections in Rat Neocortex. Cereb Cortex 25, 3818–3835 (2015).

54. E. Hay, I. Segev, Dendritic Excitability and Gain Control in Recurrent Cortical Microcircuits. Cereb Cortex 25, 3561–3571 (2015).

55. D. D. Team. (2016).

56. M. L. Hines, N. T. Carnevale, The NEURON simulation environment. Neural Comput 9, 1179–1209 (1997).

57. M. Oberlaender, A. Ramirez, R. M. Bruno, Sensory experience restructures thalamocortical axons during adulthood. Neuron 74, 648–655 (2012).

58. E. Hay, S. Hill, F. Schurmann, H. Markram, I. Segev, Models of neocortical layer 5b pyramidal cells capturing a wide range of dendritic and perisomatic active properties. PLoS Comput Biol 7, e1002107 (2011).

