## Supplemental Figures for "Thalamus drives active dendritic computations in cortex"

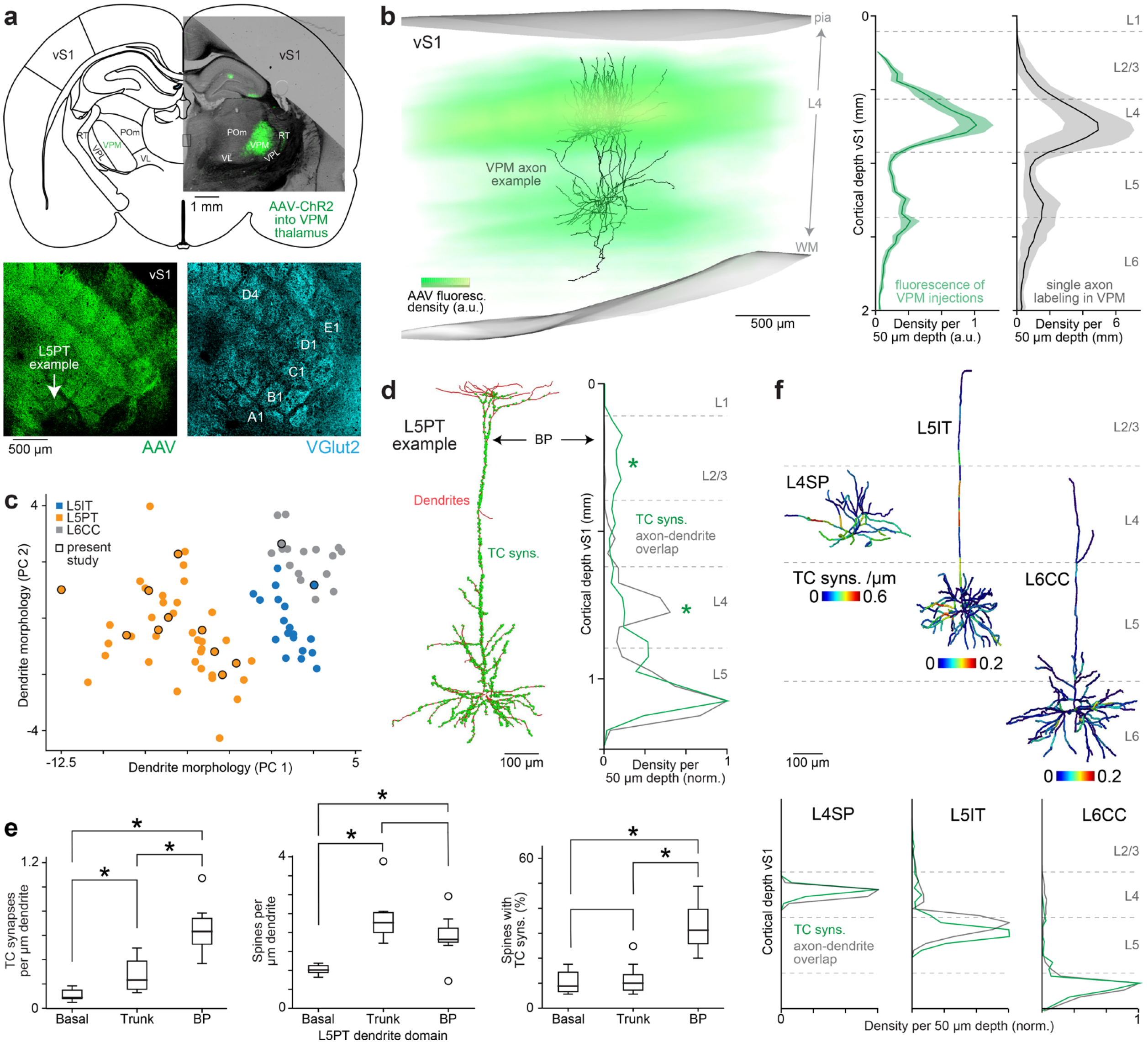


**Fig. S1 (related to Fig. 1): Control data for AAV-based mapping of TC synapses along the dendrites of *in vivo* labeled L5PTs. a.** For each animal (n=9) in which we identified putative TC synapses along the dendrites of *in vivo* recorded L5PTs (n=10), we had three criteria to confirm that AAV injections are restricted to the VPM thalamus. First, it is well known that VPM axons delineate the layer 4 barrels, whereas POm axons delineate the septa between barrels (*37*). To test this, we first cut the cortex of the AAV injected brains into consecutive 50 µm thick sections tangential to barrel cortex (i.e., vS1) and took images of the sections comprising layer 4. Bottom panels show images of layer 4 barrels for AAV (left) and VGlut2 labeling (right) for the injection site shown in the top panel. The arrow denotes the location of the trunk of the example L5PT shown in **Fig. 2-4** and **Fig. 1e** (most right). Second, we cut the remainder of the same AAV injected brains into 100 µm thick coronal sections, took images of the sections that comprised the injection site, and aligned them with the corresponding images of the Paxinos Rat Brain Atlas based on the outlines of thalamic nuclei, the hippocampus, cortex and brainstem. Top panel: image shows AAV injection site restricted to VPM thalamus corresponding to layer 4 images in the bottom panels. **b.** Third, it is well known that VPM axons terminate most densely in layer 4 and at the layer 5 to 6 border, whereas POm axons terminate most densely in layer 1 and at the layer 4 to 5 border (*37*).

To test this, we inspected the fluorescence density of AAV labeled axons across all layers of the barrel cortex. Left panel shows reconstruction of AAV fluorescence density across barrel cortex superimposed with axon reconstruction of a single relay cell in VPM thalamus (*57*). Right panels: density profiles (mean ± SD) across the layers of the barrel cortex for AAV fluorescence (green; n=9) and single VPM axons (black; n=14; modified from (*14*)). Thus, AAV injections are confined to VPM thalamus, delineate the barrels in layer 4 and show the same distribution across layers as single VPM axons. We therefore conclude that the TC synapses that we identified here along L5PT dendrites originate from VPM, not from POm thalamus. **c.** Principal components (PCs) of 21 dendritic features that discriminate between L5IT, L5PT and L6CC in rat barrel cortex (*51*). The markers without black outlines represent the dendrites of neurons that we had reported previously (*24, 35, 51*), including L5PTs (n=22) with identified long-range targets in subcortical areas (*35*). The markers with black outlines represent the neurons reconstructed in this study. Thus, the twelve neurons with somata in layer 5 for which we identified TC synapses along their dendrites represent ten L5PTs **(Fig. 1e)**, and one L5IT and L6CC, respectively (see panel f). **d.** Left panel: 3D reconstruction of the dendrites (red) and TC synapses (green) of the L5PT for which the AAV injection site is shown in panel a. Right panel: Distributions of TC synapses along the dendrites of the L5PT on the left versus axo-dendritic overlap (normalized to the respective peaks) with reconstructions of individual *in vivo* recorded VPM cells (*51*). Axo-dendritic overlap can account for the distribution of TC synapses along the basal dendrites of L5PTs (i.e., in layer 5), but predicts higher than observed densities of TC synapses for apical oblique dendrites (i.e., asterisks in layer 4), and lower than observed densities for distal apical dendrites (i.e., asterisks in layer 2/3). **e.** To demonstrate that the highest density of TC synapses around the BP of L5PTs does not reflect higher spine densities in this region, we measured spine and TC synapse densities for three dendritic domains for all reconstructed L5PTs. Left panel: the density of TC synapses around BP (0.81 ± 0.23 per µm of dendrite) is significantly higher compared to other parts along the apical trunk (0.34 ± 0.16; two-sided t-Test unpaired: $p<10^{-3}$, n=10 L5PTs), and to the basal dendrites (0.13 ± 0.05; $p<10^{-5}$). Center panel: spine densities are not significantly different around the BP compared to other parts of the apical dendrite (p=0.08), but significantly higher compared to basal dendrites ($p<10^{-3}$). Thus, while the fraction of spines that form TC synapses is largely constant across basal and apical dendrites (right panel), it is significantly higher around the BP ($p<10^{-4}$). **f.** Top panel: TC synapse densities along the dendrites of *in vivo* recorded EXC neurons in vS1 that are not L5PTs – i.e., L4SP, L5IT and L6CC. Bottom panel: axo-dendritic overlap can account for the distributions of TC synapses along the dendrites of these three example neurons.

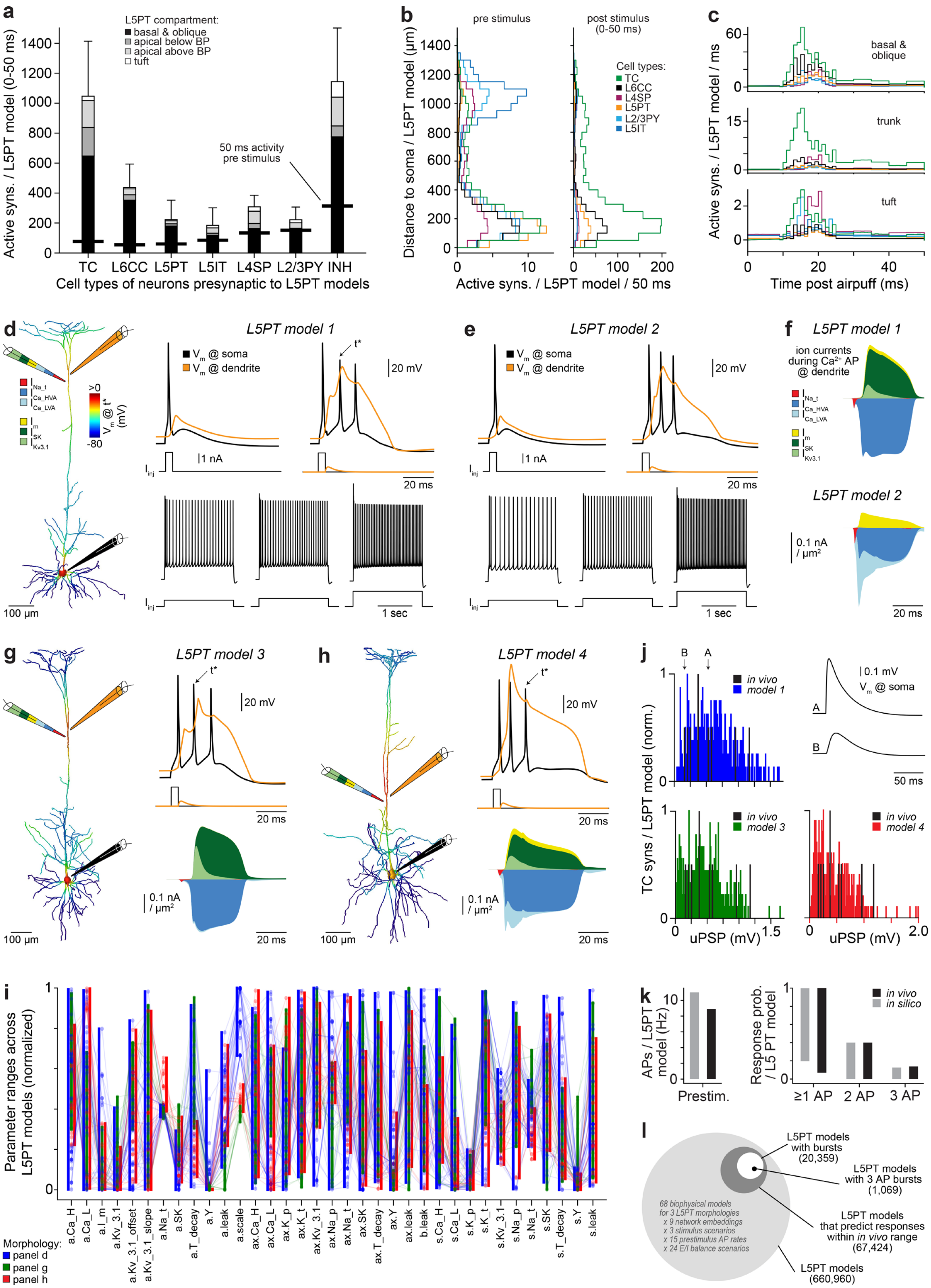

a
L5PT compartment:
basal & oblique
apical below BP
apical above BP
tuft
Active syns. / L5PT model (0-50 ms)
50 ms activity pre stimulus
TC
L6CC
L5PT
L5IT
L4SP
L2/3PY
INH
Cell types of neurons presynaptic to L5PT models
b
pre stimulus
post stimulus (0-50 ms)
Cell types:
TC
L6CC
L4SP
L5PT
L2/3PY
L5IT
Distance to soma / L5PT model (µm)
Active syns. / L5PT model / 50 ms
c
basal & oblique
trunk
tuft
Active syns. / L5PT model / ms
Time post airpuff (ms)
d
L5PT model 1
V_m @ soma
V_m @ dendrite
t*
20 mV
1 nA
20 ms
1 sec
100 µm
e
L5PT model 2
f
L5PT model 1
ion currents during Ca²⁺ AP @ dendrite
L5PT model 2
0.1 nA / µm²
20 ms
g
L5PT model 3
h
L5PT model 4
j
in vivo
model 1
model 3
model 4
0.1 mV
V_m @ soma
50 ms
TC syns / L5PT model (norm.)
uPSP (mV)
i
Parameter ranges across L5PT models (normalized)
Morphology:
panel d
panel g
panel h
k
APs / L5PT model (Hz)
Prestim.
Response prob. / L5 PT model
in vivo
in silico
≥1 AP
2 AP
3 AP
l
L5PT models with bursts (20,359)
L5PT models with 3 AP bursts (1,069)
L5PT models that predict responses within in vivo range (67,424)
L5PT models (660,960)
68 biophysical models for 3 L5PT morphologies
x 9 network embeddings
x 3 stimulus scenarios
x 15 prestimulus AP rates
x 24 E/I balance scenarios

**Fig. S2 (related to Fig. 3): Control data to account for parameter degeneracy in multi-scale models of L5PTs. a.** Quantification of spatiotemporal EXC synaptic input patterns (mean ± SD) that impinge during pre- and post-stimulus periods of 50 ms onto the dendrites of 660,960 biophysically-detailed multi-compartmental L5PT models embedded into anatomically-detailed and empirically validated network models of rat vS1 (one example for such a multi-scale model configuration is shown in **Fig. 3b-c**). The bold black lines denote the number of active synapses from each cell type during 50 ms preceding the stimulus (i.e., in anesthetized rats). The gray shadings denote the respective numbers of active synapses during 0-50 ms post stimulus (i.e., passive whisker deflection by airpuff) onto different dendritic domains as in **Fig. 1d**. Variability of spatiotemporal synaptic input patterns originates from the different dendrite morphologies of the L5PT models, their different embeddings into the network model, and the activation of their presynaptic neurons in the network model by generating Poisson spike trains based on our *in vivo* recorded firing rates for each cell type. **b.** Quantification of the same spatiotemporal synaptic input patterns as in panel a (mean), now resolved by dendritic locations of input. Note: despite generally low prestimulus firing rates during anesthesia, synapses from neurons across all layers and of all cell types imping onto L5PTs, with L5IT providing the majority of prestimulus inputs to the apical tuft dendrites in the model during these experimental conditions (left). Furthermore, even though TC synapses from VPM represent generally less than 5% of the total number of inputs that L5PTs receive, they represent the majority of active synapses after stimulus onset during these experimental conditions (right). **c.** Quantification of the same spatiotemporal synaptic input patterns as in panels b, now resolved by 1 ms time bins of input. Note: TC synapses dominate the sensory-evoked input to the trunk of L5PTs, whereas inputs to the proximal (i.e., basal and apical oblique) and distal (i.e., tuft) dendrites arise equally abundant from the IC EXC cell types. **d.** Example multi-compartmental model of the L5PT with the most superficial BP. For each multi-compartmental L5PT model, we simulated current injections into the soma (black) and/or calcium domain (orange) to capture the empirically observed dendritic (top) and perisomatic (bottom) physiology of L5PTs (*58*), including bAPs (top-left), calcium APs and burst firing when inputs to the soma and calcium domain coincide (top-right), and regular AP firing of increasing frequencies in response to sustained current injections of increasing amplitude (bottom). **e.** Second example of a multi-compartmental model for the same L5PT as in panel d. **f.** Both example models of the same L5PT morphology generate calcium APs and bursts of APs during coincidence detection equally well within the experimentally observed range, but utilize very different superpositions of ion channels to achieve these functions. We plot hyperpolarizing currents upwards: calcium-dependent (SK), fast non-inactivating (Kv3.1) and muscarinic (M) potassium channels; depolarizing currents downwards: low- and high-voltage activated calcium channels (Ca_LVA, Ca_HVA) and sodium (Na_t) channels. Currents are measured at the BP. **g-h.** Example models with diverse channel unitizations similar to those in panel f, but for different L5PT morphologies (i.e., in-between and deepest BP). **i.** Ranges of biophysical parameters across all models with acceptable dendritic and perisomatic physiology (n=68; most superficial BP: 40, in-between BP: 13, deepest BP: 15) representing the passive leak conductance and the density of Hodgkin-Huxley type ion channels on the soma (s), basal dendrite (b), apical dendrite (a), and axon initial segment (ax). Parameters are normalized to their biophysically plausible ranges (see (*58*) and **Methods**). **j.** We simulated the activation of the synapses from each presynaptic cell on the dendrites of each multi-compartmental L5PT model to optimize their respective peak conductance until their distributions matched the empirically observed uPSP amplitude distributions between different presynaptic IC cell types and L5PTs (*53*), and as shown here between cells in VPM and L5PTs (*12*). Note: the strengths of synapses hence differ between multi-compartmental models (n=68), but always meet the respective uPSP distributions observed empirically for the different TC and IC populations. **k.**

We simulated how the hence generated 660,960 multi-scale model configurations of L5PTs transform the spatiotemporal synaptic input patterns that mimic our anesthetized experimental conditions into somatic and dendritic APs. We selected those model configurations for *in silico* manipulations, training of ANNs and analyses that predicted pre- and post-stimulus activity as we had observed for these L5PTs *in vivo*. **l.** 67,424 models predicted pre- and post-stimulus activity as observed *in vivo*, of which 22,850 configurations contained sensory-evoked bursts. These models comprised configurations for all three morphologies, all nine network locations, all 68 biophysical parameter sets, all 3 whisker stimuli, all inhibitory timings, 7/8 post-stimulus EXC/INH ratios and 13/15 pre-stimulus EXC/INH ratios (see **Methods**). Thus, our simulations provide predictions for the functions of the TC→L5PT pathway that are robust despite parameter degeneracy of anatomical and functional properties of L5PTs at subcellular, cellular and network scales (within the respective empirical constraints).

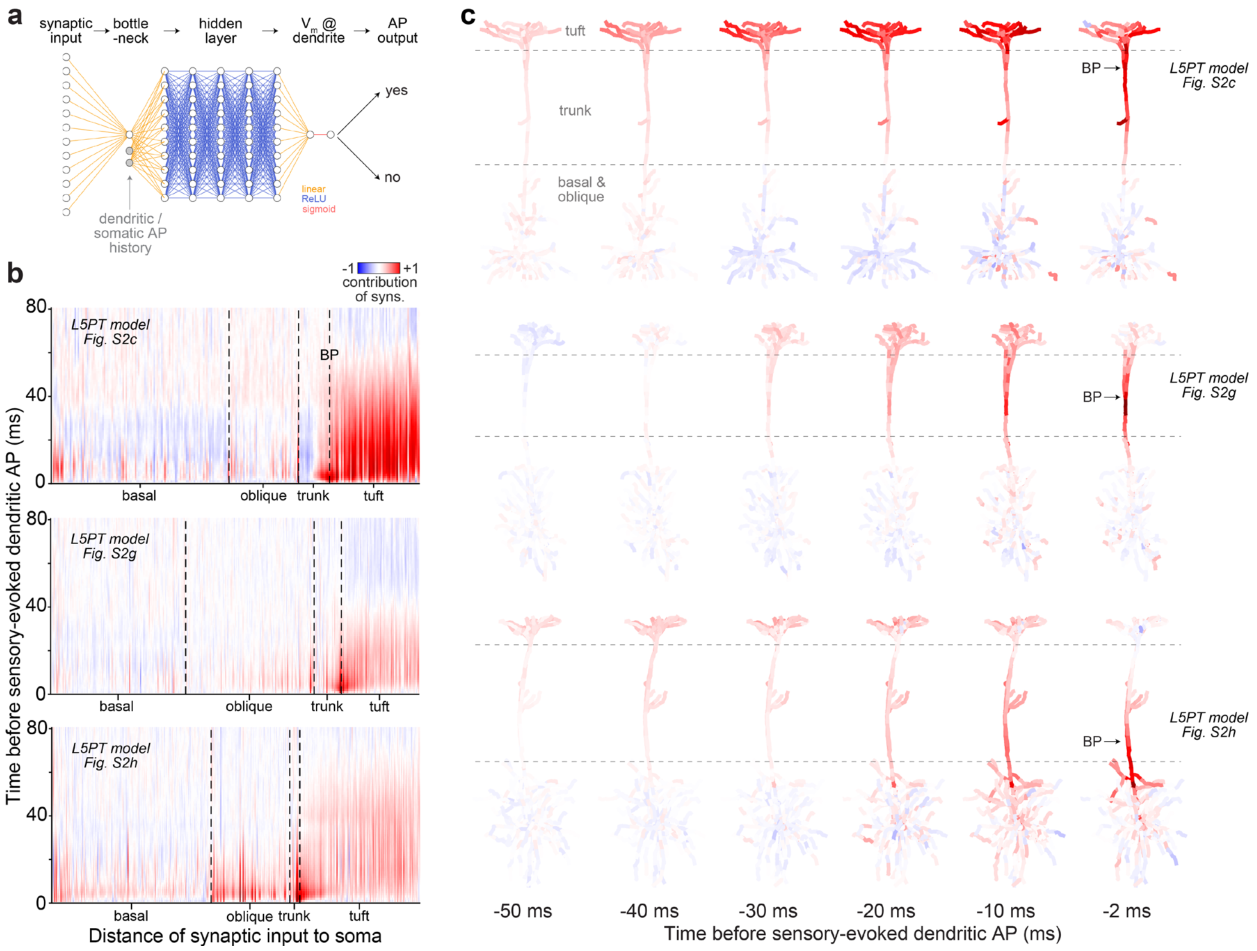


**Fig. S3 (related to Fig. 3): Control data to account for parameter degeneracy in multi-scale models of L5PTs when training ANNs. a.** We trained custom-designed ANNs on the 13,484,800 simulations of the 67,424 multi-scale model configurations that predicted pre- and post-stimulus activity as observed *in vivo* to learn general input-output relationships in L5PTs **(Movie S2)**. The ANN design is based on a multi-layer-perceptron architecture, where the input layer represents the spatiotemporal input patterns and the output layer the membrane potential at the soma or BP. The key feature in the ANN design is the low-dimensional bottleneck layer between the input and hidden ReLU layers. A dimensionality reduction of the spatiotemporal input patterns to a bottleneck dimension of 1 was sufficient to learn somatic and dendritic responses, while providing interpretable results for which locations and time points of synaptic activation contribute to which kind of somatic and dendritic responses. The time to the previous dendritic and somatic AP is concatenated to the output of layer 1. This allows the downstream layers to learn intracellular dynamics (e.g. refractory periods). **b.** Visualization of the weights for EXC synapses (INH are shown in **Movie S2**) between the input layer and bottleneck in trained ANNs that predict sensory-evoked calcium APs with high accuracy. The three examples represent multi-scale models of the three different L5PT morphologies (from top to bottom: most superficial, in-between, deepest BP). The dimension of the input layer (i.e., number of weights to bottleneck) is derived from the spatiotemporal input patterns as follows: 80 temporal 1 ms bins reflect the times of synapse activation for 80 ms preceding the time point at which the membrane potential is predicted; 238, 338 or 365 spatial ~50 µm bins (most superficial, in-between or deepest BP) reflect the dendritic locations of active synapses sorted by distance to the soma. Note: ANNs for all three L5PT morphologies and all 68 biophysical configurations reach consensus that calcium APs are driven by input to the trunk (i.e., contributions shortly before the calcium AP are highest there) and by the amount of depolarization in the tuft for periods up to ~40 ms preceding calcium APs. **c.** Visualization of the weights for EXC synapses as in panel b, mapped onto the dendrites and for different time points. Note: ANNs for all three L5PT morphologies predict that inputs around the BP shortly before the onset of calcium APs contribute the most to driving them. Thus, our predictions about the TC-coupling mechanism in L5PTs are robust despite parameter degeneracy of anatomical and functional properties at subcellular, cellular and network scales.

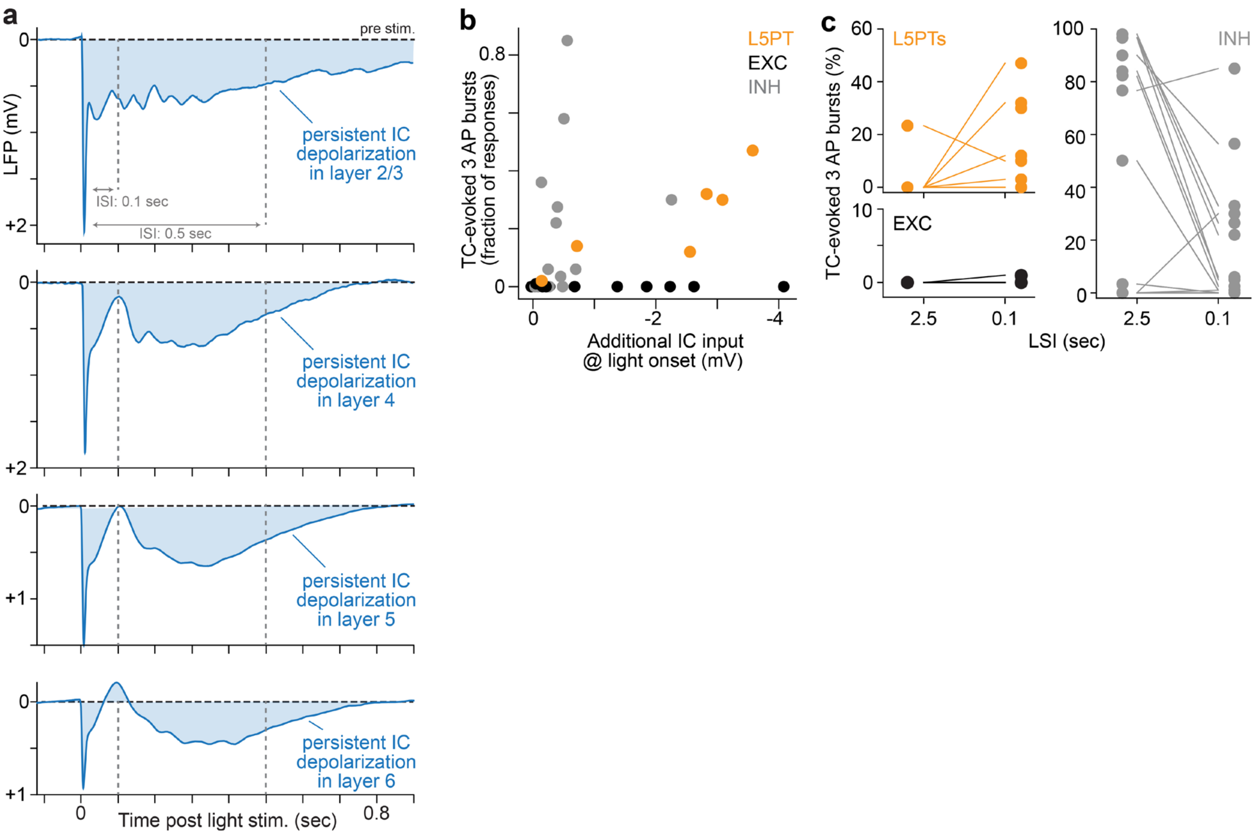


**Fig. S4 (related to Fig. 4): Control data for *in vivo* manipulations of the TC→L5PT pathway. a.** Average LFPs evoked by 10 ms light pulses analogous to that shown in **Fig. 4c**, now resolved for the different layers of vS1 (layer 2/3: n=3; layer 4: n=10; layer 5: n=44; layer 6: n=9). Note: because neurons independent of their soma depth location and cell type show elevated activity at time points up to ~0.8 seconds after light stimulation **(Fig. 4c)**, and because most of these neurons have axons that extend far beyond the layer in which their soma is located (*24*), this wave of elevated activity by IC neurons leads to a depolarized stated that persists across all layers of the barrel cortex. **b.** Light stimulated TC synapses evoke 3 AP bursts in both L5PTs and INH neurons, but not in EXC neurons of other cell types (here: L4SP, L5IT and L6CC). The fraction of trials in which L5PTs respond with 3 AP bursts increases with the amount of IC input that is additionally provided before a light pulse, which we estimated by the LFP amplitude right before the onset of each light pulse. We did not observe such a relationship for light-evoked 3 AP bursts in INH neurons. **c.** L5PTs show increased fractions of 3 AP bursts in response to light stimulations at LSIs of 0.1 seconds, which result in elevated IC activity that can persist throughout all stimulus trials **(Fig. 4f)**. In contrast to L5PTs, INH neurons show decreased fractions of light-evoked 3 AP bursts at LSIs of 0.1 seconds. Thus, the transition from TC-evoked single APs to 3 AP bursts depending on elevated IC activity is exclusively observed for L5PTs, not in other EXC and or INH neurons.
